# Effects of host sex, age and behaviour on co-infection patterns in a wild ungulate

**DOI:** 10.1101/2025.01.27.635057

**Authors:** Florian Berland, Vincent Bourret, Carole Peroz, Laurence Malandrin, Claire Bonsergent, Xavier Bailly, Sébastien Masseglia, Laurent-Xavier Nouvel, Anne-Claire Lagrée, Clotilde Rouxel, Chloé Dimeglio, Jacques Izopet, Valentin Ollivier, Thierry Boulinier, Isabelle Villena, Dominique Aubert, Vincent Sluydts, Guillaume Le Loc’h, Arnaud Bonnet, Yannick Chaval, Joël Merlet, Bruno Lourtet, Emmanuelle Gilot-Fromont, Hélène Verheyden

## Abstract

Recent zoonotic disease emergences reveal the importance of studying wildlife parasite communities. As wild hosts frequently harbour diverse parasite species, understanding the drivers of multiple infection patterns in free-ranging hosts is critical for elucidating the ecological and epidemiological dynamics of parasite communities. In this study, we analysed co-infection patterns in European roe deer (*Capreolus capreolus*) inhabiting a fragmented rural landscape in southwestern France. Using data from 130 GPS-tracked roe deer, we examined the influence of proximity to livestock, host activity levels, age, sex, and between-parasite interactions on the presence of 11 parasitic taxa. Hierarchical modelling of species communities (HMSC) revealed that proximity to livestock significantly increased the likelihood of infection with orofecally transmitted parasites (*Toxoplasma gondii*, gastrointestinal parasites). Sex and age were other key predictors, with males and juveniles exhibiting a higher frequency of parasite presence, likely influenced by hormonal and immune system differences. Activity levels showed distinct age-related effects, with higher activity levels being positively associated with increased parasite prevalence in yearlings, but not in adults. In contrast, parasite association patterns within individual hosts were weak, suggesting minimal interactions between parasite species. Our findings highlight the interplay between exposure and susceptibility in shaping co-infection patterns and underscore the value of hierarchical modelling approaches in multi-parasite systems.

**Key Findings:**

- Parasite communities, studied in 130 GPS monitored roe deer, revealed frequent co-infection.
- Proximity to livestock increased the risk of infection by parasite taxa shared with domestic animals.
- For several parasitic taxa, males and younger roe deer had a higher probability of infection.
- The effect of host activity on parasite infection was dependent on roe deer age.
- Only slight between-parasite interactions were detected within individual hosts.

## Introduction

Zoonoses are diseases transmitted between animals and humans. They account for more than 60% of current emerging infectious diseases in humans, and most of them (72%) originate from wildlife (Jones *et al*. 2008). The recent rise in emerging zoonotic diseases has underscored the importance of studying the circulation of zoonotic pathogens, and more generally parasite communities, in wildlife (Jones *et al*. 2008; Smith *et al*. 2014; Destoumieux-Garzón *et al*. 2018). Here we use the term parasite in a broad sense, including viruses, bacteria, protozoa and macroparasites. Because hosts are exposed to multiple parasitic species (Gortázar *et al*. 2016; Moutailler *et al*. 2016; de Cock *et al*. 2023), interactions may occur within the parasite community. Parasites can interact either directly when they are present simultaneously within the same individual host, or indirectly through some effects on the host that potentially influence the multiplication, shedding, and eventually transmission dynamics of other parasites (Ezenwa, 2016; Keegan *et al*. 2024). Indirect interactions can be mediated by parasite effects on the immune system that can result in increased infection intensity, enhancing the parasite transmission rate (Cattadori *al.* 2007). Some indirect interactions may also have unexpected consequences and complicate the implementation of parasite control strategies. For example, anthelmintic treatments increased the survival of African buffaloes, *Syncerus caffer*, infected with bovine tuberculosis, thereby enhancing the spread of the disease at the population scale (Ezenwa and Jolles, 2015). Other studies revealed that the treatments against nematode infections in wild mice (*Apodemus sylvaticus* and *Peromyscus* species) was associated with the suppression of competitive between-parasite interactions and resulted in an increase of protozoa infection (Knowles *et al*. 2013; Pedersen and Antonovics, 2013).

A key target of host-parasite ecological studies is thus to disentangle the effects of between-parasite interactions from other determinants of parasitism influencing the “twin pillars of infection”: exposure and susceptibility (Sweeny and Albery, 2022). Exposure represents the likelihood of an individual encountering parasites in its environment, while susceptibility describes the probability of exposure leading to infection (Viney and Graham, 2013, Civitello and Rohr, 2014).

Exposure can vary among individuals, according notably to the density and species diversity of the host community in which they are living (Arneberg *et al*. 1998; Horcajada-Sánchez *et al*. 2018), which may affect the abundance, diversity, spread and persistence of vectors and parasites. Vector and parasite abundances may notably be positively correlated to the host density, offering available resources and increasing the contact rate and the probability of transmission between hosts (Arneberg *et al*. 1998). Consequently, individuals using environments with a high density of hosts, with which they share parasites, should be more exposed to parasites and present a greater probability of infection. For example, areas of coexistence between livestock and wildlife, where host density is increased by the presence of livestock in restricted areas, may favour the exposure to different parasites shared between ungulate species (Pato *et al*. 2013; Sevila *et al*. 2014; Verheyden *et al*. 2020).

Behavioural components other than contact with other hosts can also influence exposure. It has been observed that more active individuals tend to be more often parasitized than less active ones (Dunn *et al*. 2011; Patterson and Schulte-Hostedde, 2011; Santicchia *et al*. 2019), which has been interpreted as the result of higher exposure. For example, in chipmunks, *Tamias minimus*, more exploratory individuals exhibited higher ectoparasite abundance (Bohn *et al*. 2017). Specific activities such as foraging and travelling are most often associated to exposure (Barron *et al*. 2015; Dougherty *et al*. 2018). These activities can vary between individuals in relation to their sex, age, or personality which may influence their home range size, activity pattern and propensity to use some habitats (Sih *et al*. 2004; Bonnot *et al*. 2018; Malagnino *et al*. 2021). Therefore, the inter-individual variability in habitat use and level of activity may lead to different levels of exposure to parasites (Sih *et al*. 2018; Brehm *et al*. 2024). Moreover, individuals are often co-exposed to multiple parasites, and the inter-individual variability in spatial behaviour could influence the probability of co-infection and the global structure of the within-host parasite community (Fox *et al*. 2013; Vanden Broecke *et al*. 2021, 2023).

Among other factors, host susceptibility is related to the immune system’s ability to clear parasites, which itself is influenced by individual characteristics such as age, sex, or body condition, through differences in hormone production or resource allocation (Cross *et al*. 2009). In the context of co-infection patterns, susceptibility to one parasite can also be linked to the presence of other parasites. Parasites differ in their nutritional and energetic needs and in their stimulation of host immunity (Ezenwa and Jolles, 2015; Abbate *et al*. 2024). Their co-occurrence in a single host can lead to direct or indirect, negative or positive interactions through mechanisms such as immunomodulation or competition between parasites for the host resources (Telfer *et al*. 2010; Dallas *et al*. 2019). For example, the presence of gastro-intestinal parasites may enhance the Th2 response and downregulate the Th1 response, thus increasing the susceptibility to some microparasites (Graham, 2008). This situation has been observed in African buffalo, where nematode infection induced Th1 immunity suppression, increasing susceptibility to bovine tuberculosis (Ezenwa *et al*. 2010). The relative impact of within-host parasite interactions and other determinants of immune response such as age, sex and body condition are not clearly established, however all may contribute to parasite distribution and co-infection patterns (Vanden Broecke *et al*. 2021;2023).

In order to investigate the relative significance of exposure and susceptibility factors on co-infection patterns in wildlife, we analysed the variability of parasite communities among individuals in a wild species, the European roe deer (*Capreolus capreolus*). Our aim was to disentangle the effects of various factors that may influence host exposure and susceptibility, including age, sex, behaviour and between-parasite interactions.

Roe deer populations are found throughout most of Europe, exploiting both forested and more open environments, including cultivated lands, livestock breeding areas and human infrastructures (Linnell *et al*. 2020). Such habitat use increases their likelihood for exposure to novel parasites, including those shared with humans and livestock (Sevila *et al*. 2014; Beaumelle *et al*. 2022). Additionally, there is high inter-individual variability in roe deer spatial behaviour, in terms of space use, home range structure, and activity (Bonnot *et al*. 2015, 2018; Malagnino *et al*. 2021). Moreover, immune parameters have been shown to vary among classes of age and sex and with physiological profile such as stress response in this species (Cheynel *et al*. 2017, Carbillet *et al*. 2023). These characteristics make roe deer a relevant biological model for investigating the link between exposure, susceptibility and the composition of their parasite community. Our objectives were to investigate the influence of host behaviour, age, sex and parasite interactions on the patterns of co-infection. We used data from a roe deer population living in a rural area, where individuals were regularly captured and GPS tracked between 2016 and 2022, and where the presence of 11 parasites have been assessed.

We expected several factors to influence the probability of parasite presence in roe deer.

(i) First, following the hypothesis than host density enhances parasite abundance, we predicted that exposure of roe deer to areas used by livestock and domestic animals should increase their probability to carry parasites that are shared with domestic hosts (Pato *et al*. 2013; Sevila *et al*. 2014). This may concern orofecally transmitted parasites such as Coccidia, *Nematodirus* spp. and Strongylidae (shared with livestock), Hepatitis E virus (excreted by pigs), and *Toxoplasma gondii* oocysts (excreted by cats that may live in proximity to livestock) as well as tick-borne parasites such as *Anaplasma phagocytophilum* (Chastagner *et al*. 2017).

(ii) Because immunity may differ between sexes due to hormonal differences, and may be less efficient in naïve individuals, we expected that males and young roe deer should be more susceptible to infection and thus showed higher infection probability (Cross *et al*. 2009; De La Peña *et al*. 2020). In addition, the slight differences in spatial behaviour between sex and age classes in roe deer during the reproductive period (Malagnino *et al*. 2021) could also lead to a higher exposure of males due to the greater distances travelled (Brehm *et al*. 2024).

(iii) Third, because exposure to parasites should increase with increasing level of exploratory activity, we expected individual activity level to be positively correlated with parasite presence and species richness (Bohn *et al*. 2017; Santicchia *et al*. 2019).

(iv) However, we also considered that the effects of exposure factors such as host activity may be nuanced by other host characteristics such as age or sex, which influence susceptibility and modify the likelihood of exposure leading to infection. (Sweeny and Albery, 2022).

(v) Finally, within-host interactions among parasites may modulate infection risk (Dallas *et al*. 2019). Competition between parasites exploiting the same host resources should promote negative associations in hosts, for instance among tick-borne parasites parasitizing the blood compartment, such as *A. phagocytophilum*, *Babesia* spp., *Bartonella* spp., and *Mycoplasma* spp. (Telfer *et al*. 2010). In contrast, positive associations were expected between gastrointestinal macroparasites and some microparasites through immunomodulation (Graham, 2008).

## Materials and methods

### Study site and animal monitoring

The study was conducted within the "Zone Atelier Pyrénées-Garonne", situated in the south-west of France, around the village of Saint-André (43°16′ N, 0°51′ E). This hilly terrain reaches a maximum altitude of approximately 380 m, and features a mean annual temperature of 12.3 °C and an average annual rainfall of 800 mm. The 19,000-ha site encompasses a fragmented rural habitat comprising two forested areas and more open environments, including woodlands, hedges, human dwellings, cultivated crops and pasture areas. Landscape permanent structures of the study area have been mapped, and agricultural use is monitored each year at the parcel level as part of long-term ecological studies on this site.

Livestock, including cattle, sheep, and occasionally goats and horses, graze in the pasture areas, and an indoor pig farm regularly carries out manure spreading for crop fertilisation. Pets such as cats and dogs are also observed in proximity to human dwellings. The presence of livestock was monitored weekly in each pasture during the whole grazing season in 2013 (Verheyden *et al*. 2020) (see references for methodologic details) and in part of the study area in 2021. Livestock presence changed in only 17% of the pasture area between 2013 and 2021, thus we considered that grazing practices remained constant during the study period.

Between January and March from 2016 to 2022, a total of 402 roe deer captures were performed at six distinct sites within the study area, representing 319 individuals for which 65 were recaptured at least once. Following capture, the sex was determined, and the age of each roe deer was estimated by assessing tooth condition, classifying individuals into three categories: juveniles (6 to 10 months), yearlings (18 to 22 months), and adults (older than 2 years). Samples of faeces and whole blood were taken from each captured individual for parasitological and physiological analyses, respectively. Serum was extracted after 10 minutes of centrifugation at 3000 g. Samples were refrigerated during transport to the INRAE-CEFS laboratory and stored at -20°C (blood and serum) or 4°C (faeces) until analysis.

Prior to release, roe deer were marked with ear-tags and sub-cutaneous microchips. Of the 402 captures, 307 included fitting a Global Positioning System (GPS) collar programmed to record at least one fix every 6 hours for 48 weeks. The GPS collar was equipped with activity sensors recording acceleration on X (forward/backward) and Y axis (sideways) every 5 minutes (288 records per day) (Benoit *et al*. 2020).

### Roe deer spatial behaviour

#### Home range and livestock proximity

Home range can be defined as the area used by an individual in its normal activities of food gathering, mating, and caring for the young (Burt, 1943). Roe deer home ranges were estimated using the Kernel method (Worton, 1989) with the "adehabitatHR" package in R version 4.2.3. We opted for the 95% full home range value to capture most of the area normally used by the individuals (potentially excluding unusual movements such as excursion). The GPS records started 15 days after the date of capture (between January and March) and finished 8 to 10 months later (November). As roe deer are sedentary and exhibit high spatial fidelity (Hewison *et al*. 1998), the home range observed during the GPS survey was considered to reflect the usual spatial behaviour and accurately estimate the individual’s exposure to parasite during the period preceding the capture, *i.e.,* explaining the presence of parasites at capture. However, some juvenile individuals leave their natal area and disperse to another place during the spring following the capture (Hewison *et al*. 1998). In that case, their home ranges do not represent the spatial behaviour they exhibited before the capture. Thus, we removed from the dataset juvenile individuals that dispersed during the GPS survey.

We described the presence of livestock into the home range as a qualitative variable with five levels (No livestock; Cattle; Cattle and Sheep; Cattle and Pig; Cattle and Sheep and Pig).

#### Mean daily activity

To estimate the level of roe deer activity, we used the mean daily Overall Dynamic Body Acceleration (ODBA) (Wilson *et al*. 2006). Dynamic body acceleration is deemed to be a good proxy of energy expenditure (Qasem *et al*. 2012), notably reflecting movements such as foraging and travelling (Benoit *et al*. 2020). For every individual, we summed the 288 activity measurements recorded every 5 minutes each day to calculate the total daily activity. We then averaged the daily activity over the entire monitoring period for each individual to obtain the mean daily ODBA value on the overall GPS record period.

### Parasite detection

Total genomic DNA was extracted from each whole blood sample using the NucleoSpin® Blood extraction kit (Macherey-Nagel, Düren, Germany). Negative extraction controls were performed by replacing the blood with 200 µL of sterile 1X PBS. Five parasites (*A. phagocytophilum*, *Babesia capreoli*, *Babesia venatorum*, *Bartonella* spp., *Mycoplasma* spp.) were directly detected by PCR of the genomic DNA. The presence and count of eggs were determined from the faeces for gastro-intestinal parasites (Coccidia, *Nematodirus* spp. and Strongylidae) with flotation in hypersaline solution method. For the three remaining parasites (*Borrelia* spp., *T. gondii*, hepatitis E virus), assessment of roe deer exposure was performed by antibodies detection in the sera. A recapitulation is available in Table 1 and detailed protocols for each parasite are available in the supplementary material.

**Table 1.**
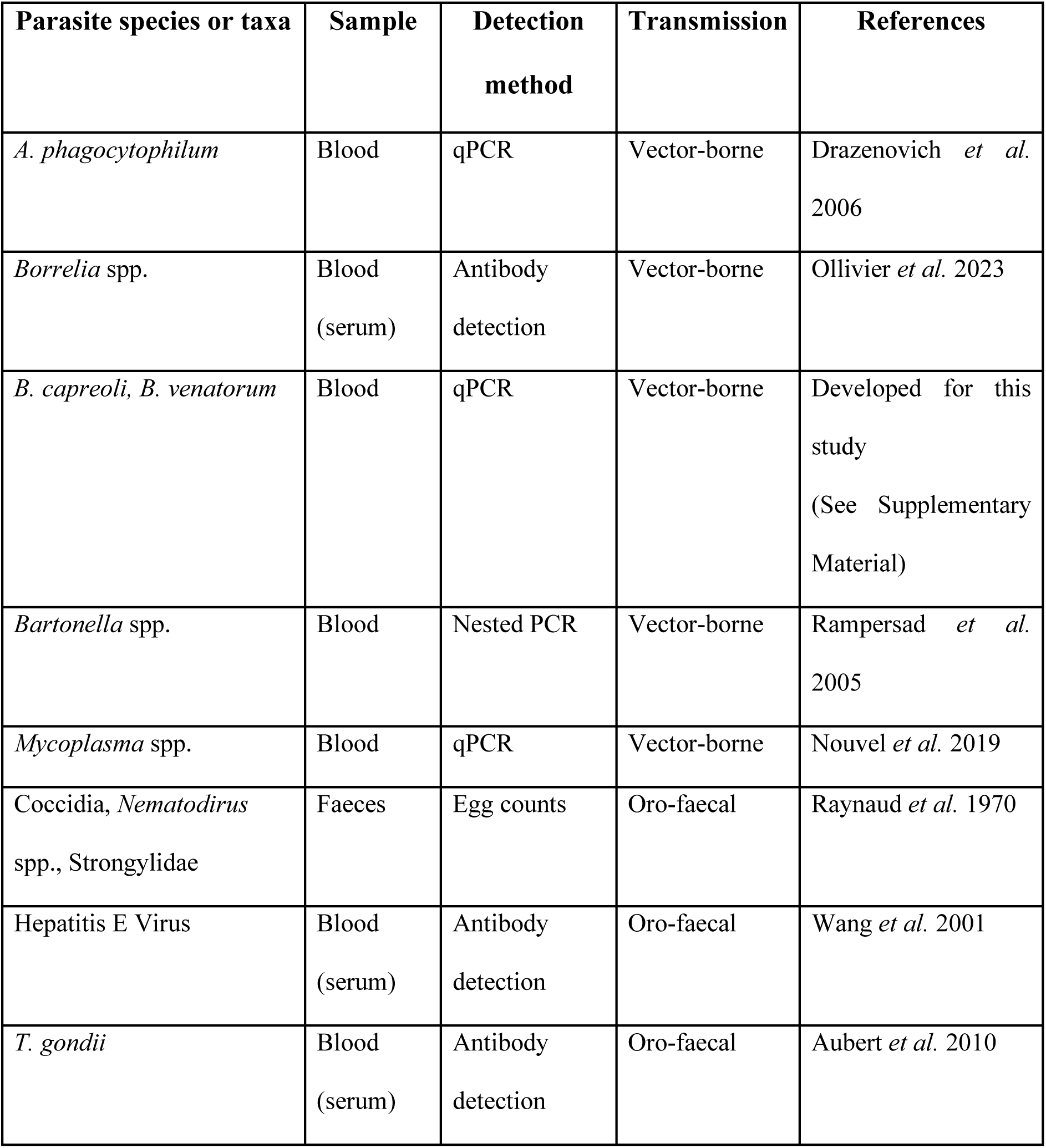
Overview of direct and antibody detection methods used for eleven roe deer parasite taxa.

### Dataset

Out of the 307 captures where GPS collar was equipped, we retained a subset of 130 captures realised between 2016 to 2017 and 2019 to 2022, representing 120 individuals, 10 of which were recaptured once. These were selected because *(i)* the animal was tested for all 11 parasites, and *(ii)* at least 75% of their home range was covered by our land-use maps.

### Statistical analysis

To investigate the influence of proximity to livestock, host activity levels, age, sex, and parasite interactions on co-infection patterns, we used Hierarchical Modelling of Species Community (HMSC), a recently developed method of hierarchical modelling (Ovaskainen *et al*. 2017). This method allows one to disentangle the different categories of factor impacting the structure of the studied parasite community (Vanden Broecke *et al*. 2021, 2023).

The HMSC is a joint species distribution model with hierarchical layers, allowing analysis of parasite responses to environmental factors, with the possibility to account for parasite phenotypic traits or phylogenetic distances (Ovaskainen *et al*. 2017). Here we considered that explanatory variables may explain either exposure (for the presence of livestock species in the home range and the level of activity) or susceptibility to parasite (for age and sex). We also included possible interactions between parasites and considered the possibility that parasites having the same transmission mode (orofecal or vector-borne) could have similar patterns.

We did not include the phylogenetic distance between the parasites due to the extreme diversity of taxa (bacteria, virus, protozoa) and the different level of identification for each parasite (family, genus, species). The response variable was the binomial presence-absence of each of the 11 parasites (or the antibodies for HEV, *Borrelia* spp. and *T. gondii*) in one individual capture event; captures of individual roe deer were considered as the sampling units. Because some individuals were captured several times and we aimed to study between-parasite interactions at the individual scale, individual identity was designated as a random effect. Additionally, we included the year of capture as a random effect, considering that individuals captured in the same year might exhibit more similar infectious profiles.

To test the effects of variables explaining exposure and susceptibility, we compared the most complete model to simpler models. The fixed effects included in the most complete comprised variables related to behaviour (livestock species in the home range, activity level, expected to determine exposure) and host characteristics, i.e., age class (juvenile, yearling or adult) and sex (male or female). As the correlation between activity and the probability of infection was expected to be shaped by host characteristics such as age and sex, we introduced the interactions between activity and both age and sex. This complete model was compared to four simpler models by removing either the interactions, exposure-related variables, susceptibility-related variables or both.

HMSC models were fitted using the R-package "Hmsc" [version 3.0-14] (Tikhonov *et al*. 2020) with default prior distributions. Posterior distributions were sampled using four Markov Chain Monte Carlo (MCMC) chains, each comprising 1,600,000 iterations, with the first 200,000 iterations discarded as burn-in and a thinning of 100 iterations. Each chain yielded 4,000 posterior samples, resulting in a total of 16,000 posterior samples. Convergence of the MCMC was assessed using the R-package "ggmcmc" [version 1.5.1.1] (Fernández-i-Marín, 2016), which estimated potential scale reduction factors (Ovaskainen and Abrego, 2020) for model parameters and manual parameter inspection using graphical tools included in the package.

Explanatory and predictive powers were both assessed using the AUC (Pearce and Ferrier, 2000) and Tjur’s R² (Tjur, 2009) values. Explanatory power was estimated as the proportion of variance in the complete dataset explained by the model parameter average estimates. We then split the dataset randomly into three thirds, and calculated the proportion of variance explained by the same global parameter estimates in each of the three randomly generated datasets. The average of these three proportions was the model predictive power.

Model selection was based on the value of the WAIC (Watanabe, 2010) and of the other parameters described previously (Tjur’s R² and AUC). The quality of the model was defined by a low value of WAIC and high value of Tjur’s R² and AUC.

## Results

### Infection and co-infection frequencies

All 130 roe deer were infected with or had antibodies against at least one of the eleven parasites analysed. Additionally, 129 of them (99.2%) had been exposed to at least two parasites. The most frequent number of parasite taxa detected in a host was four (found in 35 individuals, 26.2%), and the maximum number of parasite taxa, found in two individuals, was eight (Fig. 1).

**Figure 1.**
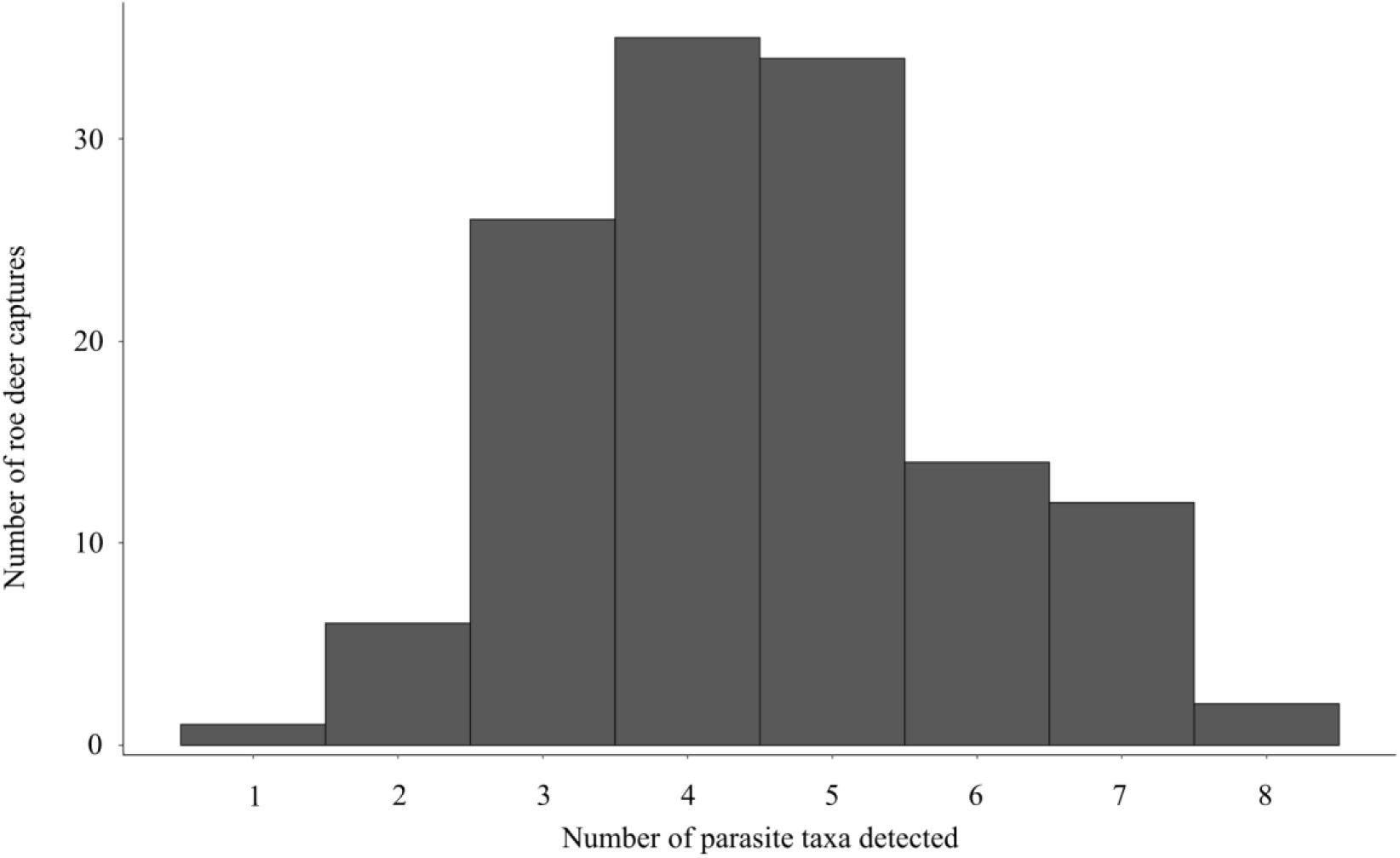
Parasite taxa richness distribution in the studied roe deer population (N = 130).

Infection frequencies varied significantly between parasites (Fig. 2). For vector-borne parasites, two taxa were highly present: *A. phagocytophilum* and *B. capreoli*, infecting 95.4% and 90% of individuals respectively. Infection frequencies were lower for other vector-borne parasites, with 30% for *Bartonella* spp., 19.2% for *B. venatorum*, 17.7% for *Borrelia* spp., and 13.8% for *Mycoplasma* spp. (Hemoplasma like).

**Figure 2.**
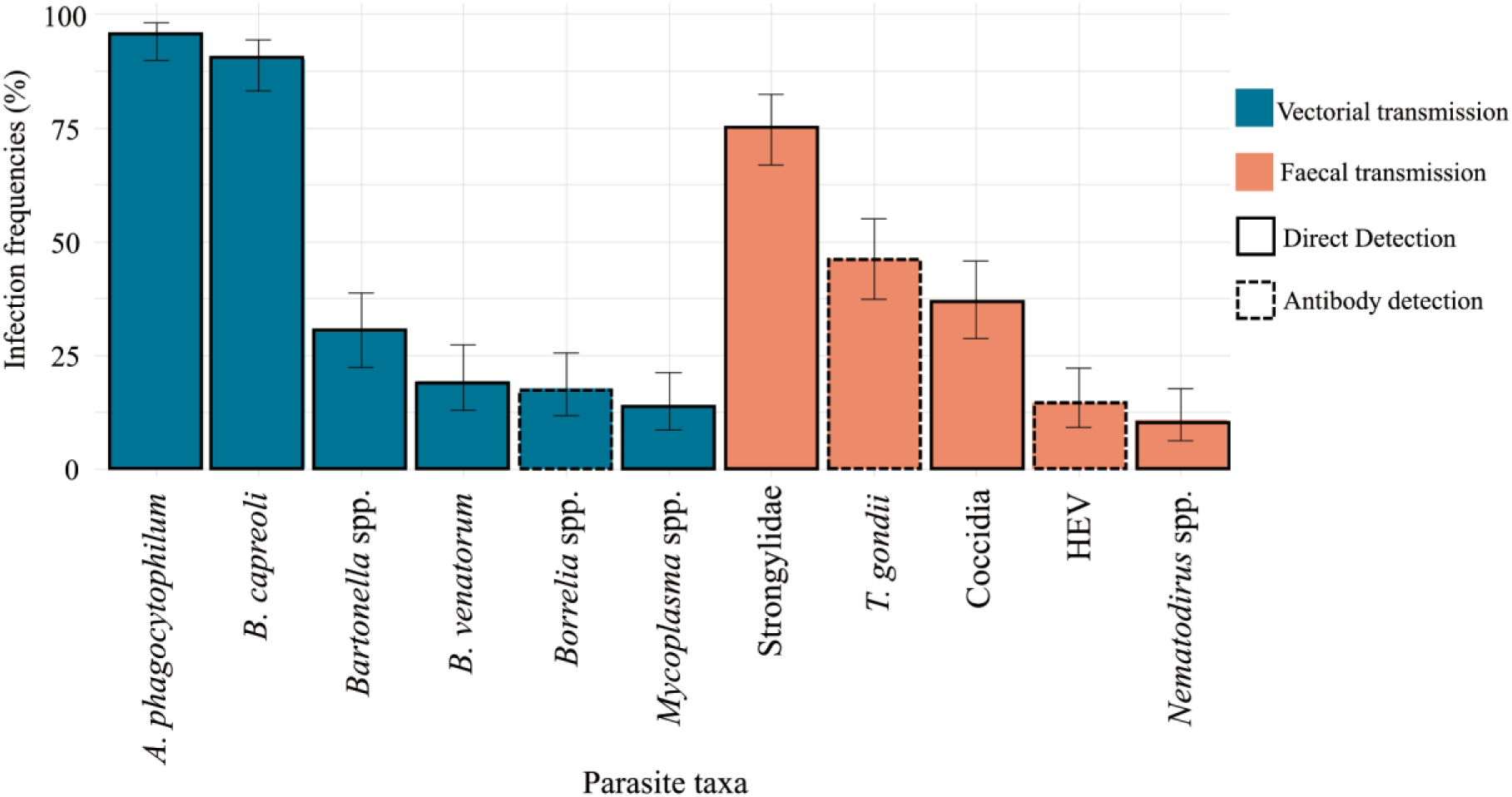
Infection frequencies for 11 parasites in 130 roe deer sampled from 2016 to 2017 and 2019 to 2022 in South-West France.

Among parasites transmitted through faecal contamination, Strongylidae exhibited the highest infection rate, with 75.4% of individuals infected. Infection rates for other taxa were lower, with 46.2% for *T. gondii*, 36.9% for Coccidia, 14.6% for *Hepatitis E Virus*, and 10.8% for *Nematodirus* spp.

### Model selection, HMSC convergence and fit

Following model comparison, we kept the most complete model including exposure and susceptibility variables together with interactions between age and activity, and sex and activity. This model had the highest explanatory and predictive power with the second-lowest WAIC value (Table 2).

**Table 2.** Model characteristics and performance to account for multiple infection patterns in 130 wild roe deer.

| Model | Random Effects | Fixed Effects | Explanatory mean Tjur R<sup>2</sup> | Explanatory mean AUC | Predictive mean Tjur R<sup>2</sup> | WAIC |
| --- | --- | --- | --- | --- | --- | --- |
| Constant | No | No | -0.0001 | 0.480 | -0.006 | 5.230 |
| Exposure | Year + Individual | Livestock + Activity | 0.107 | 0.814 | 0.019 | 5.061 |
| Susceptibility | Year + Individual | Age + Sex | 0.101 | 0.821 | 0.037 | 4.925 |
| Exposure & Susceptibility no interaction | Year + Individual | Livestock + Activity + Age + Sex | 0.132 | 0.839 | 0.039 | 5.015 |
| Exposure & Susceptibility with interaction | Year + Individual | Livestock + Activity*Age + Activity*Sex | 0.155 | 0.854 | 0.041 | 4.994 |

Visual inspection and potential scale reduction factor evaluation for β, Ɣ and Ω parameters, representing respectively host related variables effect, parasite traits and host related variables association, and parasite associations, confirmed the convergence of the HMSC model. The averaged potential scale reduction factors were estimated to 1.0004 (max = 1.004) for β parameters, 1.0004 (max = 1.003) for the Ɣ parameters, and 1.0016 (max = 1.0117) for the Ω parameters.

The explanatory power of the selected model was satisfactory, with a mean AUC of 0.850 (range: 0.731 - 0.970) and a mean Tjur R² of 0.155 (range: 0.070 - 0.288) meaning that the model captures an average difference of 15% in predicted probabilities between individuals infected and uninfected by each pathogen. This moderate explanatory power shows that the model includes explanatory variables that are relevant for partially understanding infection factors. The predictive power was lower, with an average AUC value of 0.56 (range: 0.29 - 0.77) and an average Tjur R² value of 0.041 (range: -0.02 - 0.09). These values nonetheless indicate a good model fit with the data used.

### Livestock community in the home range

The mean home range size of roe deer was 246.6 ha, varying from 5 to 1,987 ha. Most home ranges contained pasture used by cattle (93.1%) and some contained sheep pasture (29.2%), or fields spread with pig manure (30.0%) issued from a single pig farm. Some roe deer were exposed to two species (cattle and pigs for 19.2%, cattle and sheep for 18.5%) or to the three domestic species (10.8%).

The presence of livestock species in the home range accounted for 12.8% (Fig. 3) of the inter-individual variation in roe deer parasite community. Roe deer living in home ranges containing the three domestic species - cattle, sheep, and pig - displayed significantly higher prevalences of *T. gondii* (mean β = 0.93, 95% CI: 0.23-1.70) and Strongylidae (mean β = 0.75, 95% CI: 0.06-1.47), and tended to be more often infected by Coccidia than roe deer not exposed to these animals. The latter trend also held for roe deer exposed to cattle and pigs (Fig. 4, A). A positive tendency was also revealed between the overall probability of infection by oro-faecal parasites and the presence of the three livestock species (Fig. S2). In addition, roe deer exposed only to cattle tended to be more often infected by *B. capreoli* and *A. phagocytophilum* and less often infected by *Bartonella* spp. Roe deer exposed to cattle and sheep were significantly less often infected by *Bartonella* than roe deer not exposed to any livestock species (mean β = -0.61, 95% CI: -1.28 - -0.01).

**Figure 3.**
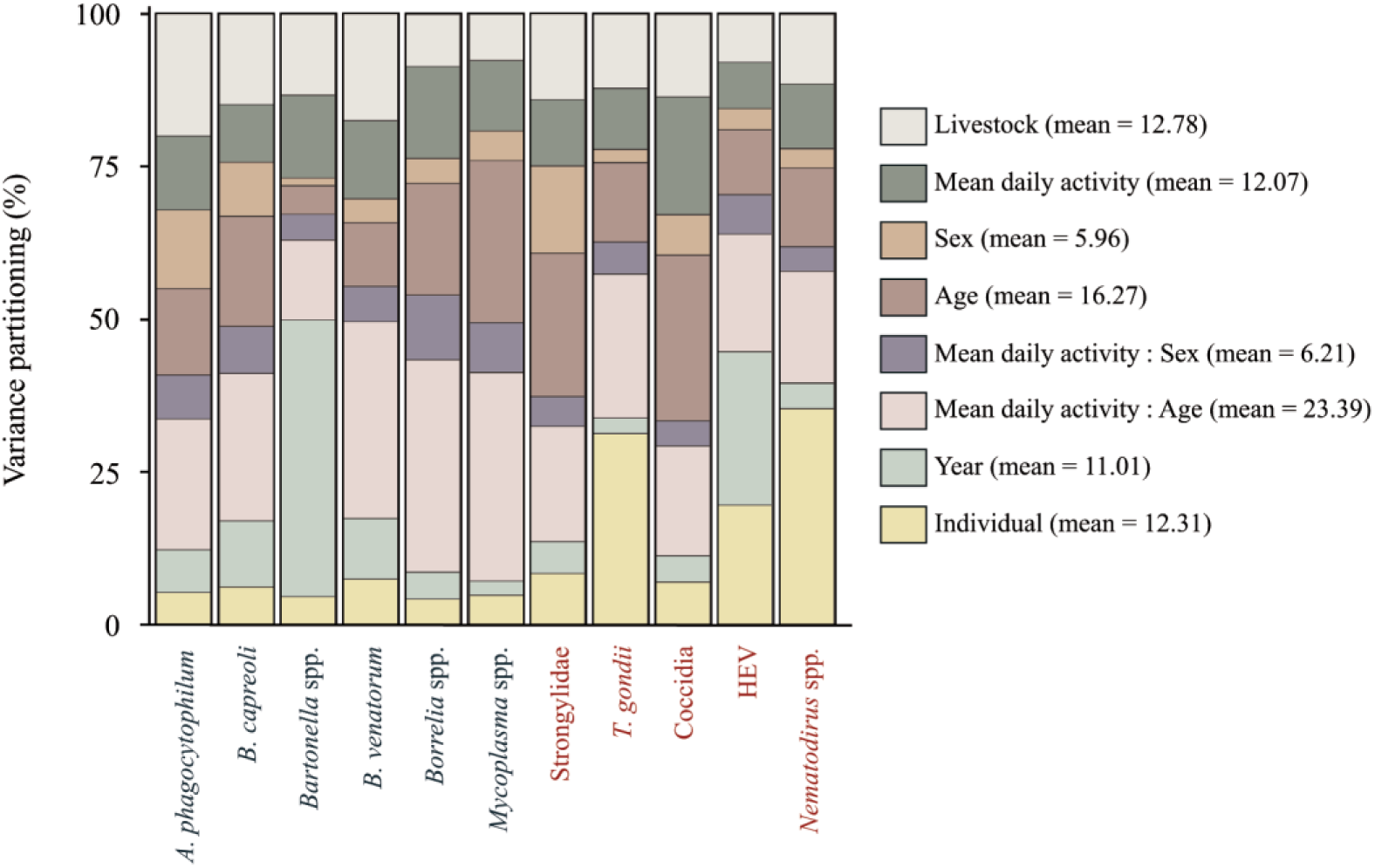
Variance partitioning among the fixed and random (year and individual) effects from the HMSC. The height of the bar represents the relative proportion of variance explained (measured with Tjur R²) achieved by each variable for the different parasite species. The legend gives the mean variance proportions for each fixed and random effect within the model, averaged over the different parasites. Dark blue coloured parasite names correspond to vector-borne parasites whereas red coloured parasite names correspond to oro-faecal transmitted parasites.

**Figure 4.**
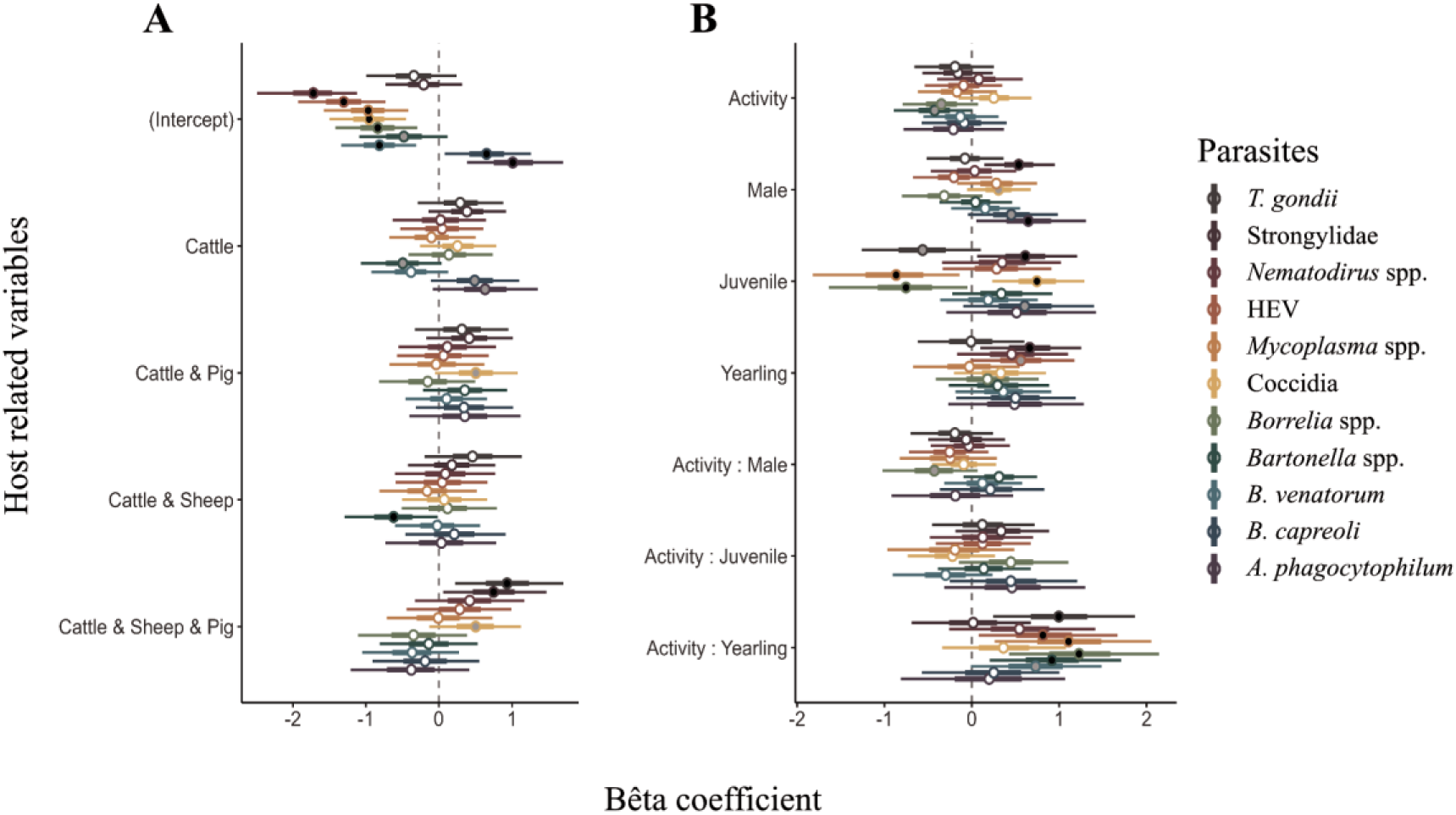
Estimation of beta coefficients for livestock presence (**A**) and host characteristics (**B**) on the probability of infection for the 11 tested parasites. The circle on each bar represents the mean of the beta coefficient, and the bar the 95% confidence interval of its values distribution. A black circle means the effect is statistically significant (95% of the beta parameter distribution is > or < to 0), a grey circle means the effect is a tendency (between 90% and 95% of the beta parameter distribution is > or < to 0) and a white means a non-significant effect.

Regarding predicted parasite taxa richness using β parameters, individuals without livestock in their home range tend to have a lower mean parasite richness (Mean Parasite Richness (PR) = 3.97, CI: 2.96-5.11) compared to those with at least one livestock species, such as cattle (Mean PR = 4.26, CI: 3.57-5.12), cattle and sheep (Mean PR = 4.09, CI: 3.21-5.09), cattle and pig (Mean PR = 4.89, CI: 4.08-5.86), or all three species (Mean PR = 4.81, CI: 3.82-5.94) (Fig. S1). The highest parasite taxa richness occurred in home ranges where cattle and pigs were present.

### Age and sex

Roe deer sex and age accounted for 5.96% and 16.27% (Fig. 3) of the variation of parasite presence, respectively. Males had a significantly higher infection rate of *A. phagocytophilum* (mean β = 0.64, 95% CI: 0.05-1.30) and Strongylidae (mean β = 0.64, 95% CI: 0.05-1.30), and tended to show a higher frequency of infection with Coccidia and *B. capreoli* than females (Fig. 4, B).

Among the three defined age categories, juveniles were significantly more likely to be infected with gastrointestinal parasites such as Strongylidae (mean β = 0.61, 95% CI: 0.06-1.20) and Coccidia (mean β = 0.74, 95% CI: 0.23-1.29), and tended to be more frequently infected with *B. capreoli* than adults. In addition, juveniles were less often seropositive to *Borrelia* spp. (mean β = -0.75, 95% CI: -1.64 - -0.05) and less often infected by *Mycoplasma* spp. (mean β = -0.86, 95% CI: -1.82 - -0.13), and tended to be less often seropositive to *T. gondii* compared to adults. Yearlings also had a higher probability of infection with Strongylidae and a tendency for a higher prevalence of HEV compared to adults. They also tended to be globally more infected by oro-faecal parasites (Fig. S2).

### Roe deer activity

The level of activity ODBA varied between a mean daily value of 1,702 to 38,677 in our study data. Activity and the interactions between activity and sex and activity and age accounted for 12.07%, 6.21%, and 23.39% (Fig. 3) of the variation in parasite infection, respectively. The positive slope of the relationships between activity and infection probability were significantly stronger for yearlings (compared to adults) for *T. gondii* (mean β = 0.99, 95% CI: 0.24-1.86), HEV (mean β = 0.81, 95% CI: 0.07-1.66), *Mycoplasma* spp. (mean β = 1.10, 95% CI: 0.25-2.05), *Borrelia* spp. (mean β = 1.23, 95% CI: 0.42-2.14), and *Bartonella* spp. (mean β = 0.92, 95% CI: 0.20-1.71) and tended to be stronger for *B. venatorum* (Fig. 3, B). This effect of yearling activity was also significantly positively correlated with the overall probability of infection by vector-borne parasites and tended to be positively correlated with the probability of infection by oro-faecal parasites (Fig. S2).

This age-dependent effect of activity was also reflected in parasite taxa richness, with a significant positive correlation between activity and parasite richness for yearlings (posterior support value = 0.99) (Fig. 5, A). In contrast, in adults, parasite taxa richness tended to decrease as host activity increased (posterior support value = 0.89) (Fig. 5, B).

**Figure 5.**
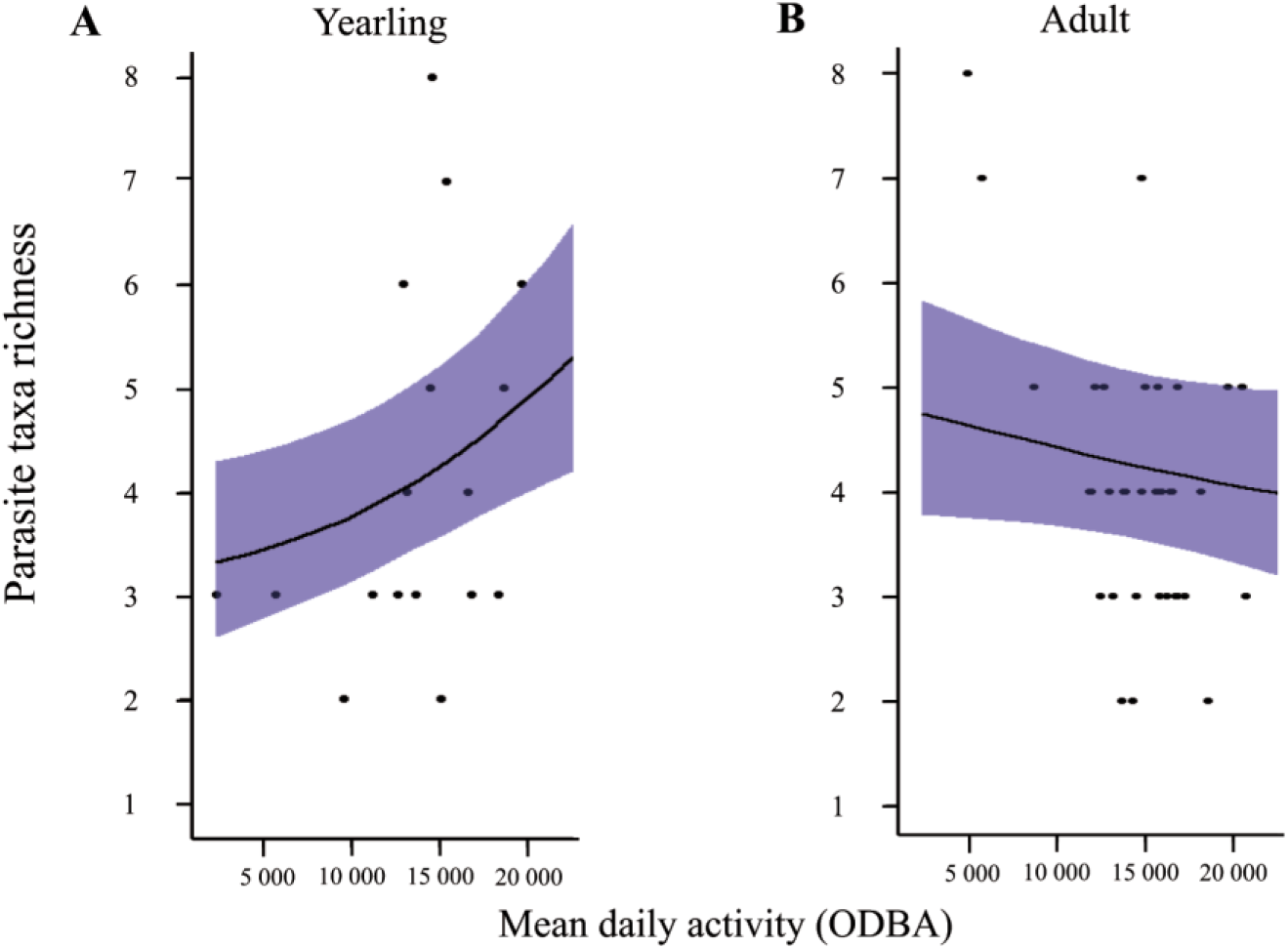
Relationship between host activity and parasite species richness predicted by the HMSC model for yearling and adult roe deer. Sex and presence of livestock species in the home range have been fixed as the more probable values in the dataset (female and cattle). Dots represent observed data; the line represents the predicted mean parasite richness for every activity value and the blue area represents the 95% confidence interval of the estimate.

### Parasite association

Residual parasite co-occurrence after accounting for exposure to domestic animals, activity, age and sex effects, was estimated at the year and individual random effect levels.

At the year scale, a significant positive co-occurrence was found between HEV and *Bartonella* spp., indicating that the HEV seroprevalence each year was highly correlated with *Bartonella* infection rates in the same year (Fig. 6, A). Additionally, *B. capreoli* showed slight tendency to co-occur with *A. phagocytophilum*, *B. venatorum* and *Borrelia* spp. antibodies. We also found a tendency for a negative co-occurrence of both *Bartonella* spp. and HEV with *A. phagocytophilum*, *B. capreoli*, *B. venatorum*, *Borrelia* spp., and with Strongylida also for *Bartonella* spp. only.

**Figure 6.**
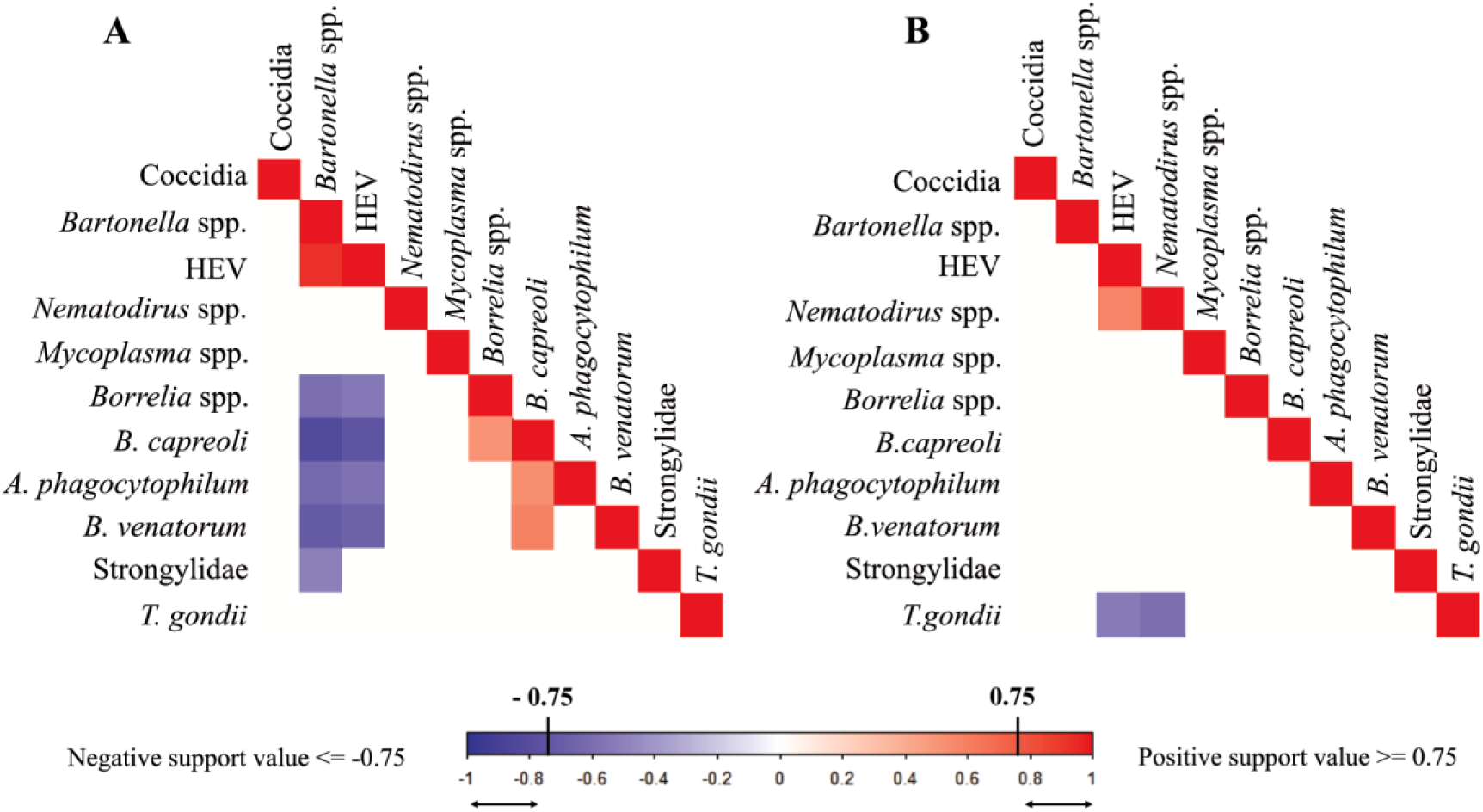
Associations between parasite presences in roe deer, after considering the effects of spatial behaviour, activity and age, sex related effects, at the year level (**A**) or at the individual level (**B**). Blue indicates a negative association whereas red indicates a positive association. Colours are shown only for a |posterior support value| > 0.75.

At the individual scale, only tendencies of negative associations between *T. gondii* antibodies and *Nematodirus* spp. and between antibodies to *T. gondii* and HEV were observed, whereas HEV antibodies and *Nematodirus* spp. tended to be positively associated (Fig. 6, B).

## Discussion

As expected, nearly all roe deer were co-infected by several parasites with most often three to five parasites among the 11 considered. This result emphasises the relevance of considering parasite communities and potential parasite-parasite interactions when studying infection patterns (Telfer *et al*. 2010). The HMSC analysis allowed us to disentangle the effects of these factors with a satisfactory fit despite the complexity inherent to data recorded in a wildlife population living in a complex environment (Vanden Broecke *et al*. 2021, 2023). As expected, exposure related factors - the presence of livestock and the level of activity - and susceptibility related factors - age, sex and between-parasite interactions - both influenced the co-infection patterns in roe deer.

### Infection patterns in relation to roe deer exposure to domestic ungulates

Consistent with our first prediction, exposure of roe deer to areas used by several livestock species increased their likelihood of harbouring parasites shared with domestic animals. Compared to unexposed individuals, roe deer exposed to areas used by cattle, pig (manure) and sheep exhibited a higher prevalence of Strongylidae and *T. gondii*. A tendency was also detected for Coccidia, which also held in the absence of sheep. These results did not hold with any other livestock species combination. This is likely due to unequal domestic species occurrence across the study area. Pigs (and pig manure) and, to a lesser extent sheep, used a limited space in our study area, whereas cattle are more widely distributed, with most roe deer exposed to them. A quantitative analysis considering the abundance of domestic animals and the intensity of roe deer infection, rather than simple occurrences of domestic animals and parasites, could have revealed a “burden intensity effect” of roe deer exposure to cattle as shown for parasitic helminths in the same area (Verheyden *et al*. 2020). In addition, because the various gastrointestinal parasite species were not easily distinguishable using faecal egg counts, we could not detect more specific host related effects of exposure, even though sheep and roe deer are known to share several gastrointestinal parasite species in this area such as *Haemonchus contortus* (Beaumelle *et al*. 2022).

Pigs are known to be infected by *T. gondii* (Lehmann *et al*. 2003; Djokic *et al*. 2016) and Coccidia (Delsart *et al*. 2022). However, these parasites are rarely (Coccidia) or never (*T. gondii*) excreted by pigs, reducing the likelihood of interspecific transmission from pig to roe deer. Nevertheless, compared to roe deer not exposed to domestic ungulates, individuals exposed to cattle, sheep, and pig in their home range are likely to live closer to farms and livestock breeding infrastructures where cats can be abundant. Indirect transmission of cat-derived *T. gondii* oocysts to roe deer, through cat encroachment into roe deer habitat or manure dispersion, could thus explain the significant positive correlation between the occurrence of three domestic ungulate species and the infection by *T. gondii*, as noted in other wild species (Gotteland *et al*. 2014; Barros *et al*. 2018).

Unexpectedly, roe deer exposed to cattle and sheep exhibit a significant lower prevalence of *Bartonella* spp. Roe deer and cattle can both be infected by *Bartonella bovis* (Skotarczak and Adamska, 2005; Cherry *et al*. 2009) and we could expect a higher probability of infection for individuals using cattle pastures. However, livestock grazing has been shown to reduce the density of rodents and the abundance of fleas (McCauley *et al*. 2008; Bueno *et al*. 2012) which are the main vector of *Bartonella* spp. (Billeter *et al*. 2008). Thus, the negative correlation found in our model could come from a negative effect of livestock grazing on the abundance of *Bartonella* spp. vectors reducing the likelihood of infection for roe deer using these areas.

Overall, our analysis suggests that parasite richness in roe deer tended to increase with the number of domestic ungulate species they were exposed to. This could be seen as consistent with the “host-diversity begets parasite-diversity” concept (Johnson *et al*. 2016). However, in addition to the introduction of new parasite species by livestock into the environment, the presence of livestock could affect the probability of roe deer infection by increasing the density of competent hosts and related parasites in the area. Thus, it could increase the roe deer exposure to several parasites and the average parasite richness of co-infections.

### Infection patterns in relation to age and sex

In many mammal species, parasitism has been shown as male-biased (Perkins *et al*. 2003; Ferrari *et al*. 2004). These sex-based differences in infection probability are often associated to susceptibility factors such as the immunosuppressive effect of testosterone (Klein, 2000; Foo *et al*. 2017; De La Peña *et al*. 2020) or males reduced investment in immunity compared to females (Rolff, 2002) but they also can be associated with exposure factors that can differ depending on the sex such as home range size, activity, or sociability (Brehm *et al*. 2024). Here, male individuals were significantly more often infected by *A. phagocytophilum* and Strongylidae, and tended to be more often infected by Coccidia and *Bartonella* spp. Testosterone is regarded as an immunosuppressive factor that increases the overall probability of infection. The production of androgen hormones may notably reduce the Th2 response (González *et al*. 2010), which is normally upregulated against gastrointestinal parasite infections (Maizels *et al*. 2004) and so increase the probability of infection by parasites such as Strongylidae. Concerning the tick-borne parasites, a higher prevalence and intensity of infection have also been found for *Anaplasma marginale* in African buffalo males (Sisson *et al*. 2023). The authors invoke possible differences in physiology and immunity leading to a higher susceptibility of males compared to females, as shown in a range of mammalian species (Foo *et al*. 2017). This sex-based difference in tick-borne parasite prevalences can also be due to morphological differences. Being on average 11% heavier than females (Chirichella *et al*. 2023), male roe deer could host more ticks than females (Kiffner *et al*. 2011) increasing their exposure to tick-borne parasites. Moreover, males are known to be more active during the reproductive period (Malagnino *et al*. 2021), which may increase their likelihood of encountering vectors and parasites and thus their probability of infection (Brehm *et al*. 2024). Concerning the effect of age, young individuals may not yet have developed acquired immunity, making them more vulnerable to infections requiring a specific immune response (Gasparoni *et al*. 2003; Body *et al*. 2011). In our study, younger individuals were more likely infected by Strongylidae (juveniles and yearlings) and Coccidia (juveniles). The Th2 immune response is generally deemed to be effective against helminths (Maizels *et al*. 2004), and age-related shifts from Th1 to Th2 immunity have been revealed in other mammals (Gardner and Murasko, 2002; Kovacs *et al*. 2004). Such a shift could also occur in roe deer, causing helminth infection probability to decrease over the growth phase age and the immunity development.

Juveniles were also less infected by two vector borne parasites: *Mycoplasma* spp. and *Borrelia* spp. These differences can reflect ticks being more likely to attach to adults due to their larger size (Vor *et al*. 2010; Kiffner *et al*. 2011), increasing their exposure to vectorized pathogens compared to juveniles. Moreover, the smaller size of juveniles and their activity, which starts after the peak of nymph abundance in spring, may make them more exposed to larvae, which take their first blood meal (Kahl and Gray, 2023), reducing their probability of infection by parasite without transovarian transmission such as *Borrelia* spp. and *Mycoplasma* spp (Zhioua *et al*. 1994; Obiegala *et al*. 2017).

### Infection patterns in relation to activity

As expected from previous studies (Bohn *et al*. 2017; Santicchia *et al*. 2019), the prevalence of several parasites (*T. gondii*, HEV, *Mycoplasma* spp., *Bartonella* spp., *Borrelia* spp., *B. venatorum*) significantly increased with roe deer activity level, but only in yearlings. Parasite taxa richness also increased with activity in yearlings and this correlation tends to reverse at the adult stage.

In our data, a significant proportion of yearlings had undergone postnatal dispersal during the previous year (34% of individuals in our population) (Debeffe *et al*. 2012). This event is stressful and resource-consuming, potentially impacting yearling immunity and susceptibility (Bonte *et al*. 2012). Moreover, this dispersal exposed yearlings to new environments and parasites, to which they may be more susceptible (Teitelbaum *et al*. 2018). Consequently, the impact of activity on exposure could be likely more significant for yearlings, leading to a greater number of infections.

However, the opposite correlation between activity and parasite richness found for adults could come from the development of the acquired immunity in the adult stage in consequence of previous exposures (Sweeny and Albery, 2022). The individual repeatability of movement and space use have been highlighted in the studied roe deer population (Gervais *et al*. 2020). More active adults were probably also more active yearlings which were more exposed to parasites during this life stage. As a result, more active adults may have developed a greater and more diversified acquired immunity, making them more resistant to infection (Hawley *et al*. 2011, Sweeny and Albery, 2022) and reversing the correlation between activity and co-infection. Complementary mechanisms may be involved: the higher activity in adults may reflect a better body condition and so, a higher resistance to infection (Goosens *et al*. 2020), or reduced adult roe deer activity may be a consequence of parasitism (Hofmann-Lehmann *et al*. 2004; Poulin, 2013; McArdle *et al*. 2018).

This age-dependent effect of activity highlights the necessity of considering both exposure and susceptibility factors and their interactions. Exposure and susceptibility can vary independently or synergistically, depending on numerous ecological factors, which complicate their inference on the probability of infection (Sweeny and Albery, 2022).

### Parasite associations and co-infection patterns

We hypothesized that parasite co-infection patterns might partly result from immunomodulation or resource competition between parasites within the host. However, only a few notable associations were observed (Fig. 6).

*Bartonella* spp. and HEV antibodies showed a strong positive association at the yearly level, meaning that the presence of one parasite in the roe deer population during a given year was significantly and positively associated with the presence of the other. Moreover, both were negatively associated with tick-borne parasites such as *Anaplasma phagocytophilum*, *Babesia* species, *Borrelia* spp., and orofecally transmitted Strongylidae. Conversely, tick-borne parasites exhibited positive associations among themselves, suggesting a division between a group of tick-borne parasites and a group comprising *Bartonella* spp. and HEV. The positive association between *Bartonella* spp. and HEV was not due to the few individuals captured in 2019 (N=7), all of which were negative for both parasites, as this trend persisted even when these individuals were excluded from the dataset. This underscores the potential role of year-specific environmental factors, such as temperature, humidity and vegetation, in shaping parasite or vector survival (Storey and Phillips, 1985; Waruiru *et al*., 2000; Gubler *et al*., 2001; Turner and Getz, 2010). For example, *Bartonella* spp. and HEV may benefit from increased annual rainfall and humidity, which could boost flea abundance (Gracia *et al*. 2012), which are the primary vectors of *Bartonella* spp., and increase water flow, facilitating HEV transmission (Tricou *et al*. 2020). Additionally, recent studies have highlighted competition among ectoparasites (Lutermann *et al*., 2015). Tick infestations, for instance, have been shown to compete with the occurrence of lice (Hoffmann *et al*., 2016) and fleas (Krasnov *et al*., 2010). Consequently, environmental conditions favouring flea proliferation may enhance the transmission of *Bartonella spp.* and HEV outbreaks while reducing tick infestations and the transmission of tick-borne parasites, potentially explaining the patterns observed in our results. At the individual scale, we did not observe the negative association expected between blood parasites sharing the same resources. We only detected some tendencies of negative association of *T. gondii* with both HEV and *Nematodirus* spp., whereas the two later parasites tended to be positively associated. Although some studies suggest that the immunosuppressive effects of hepatitis viruses may enhance helminth infections (Cox, 2001), and that *T. gondii* immunomodulation could reduce microparasite infections (Ahmed *et al*., 2017), interpreting these results with certainty remains challenging. The slight tendencies observed at the individual level indicate that between-parasite interactions in roe deer are likely negligible compared to other factors influencing exposure and susceptibility. Furthermore, these findings may be limited by the dataset, which includes few recaptures and lacks longitudinal monitoring of individuals, thereby restricting the ability to observe changes in parasite communities over time.

## Conclusion

This study highlights the value of integrating exposure and susceptibility factors, along with their interactions, to better understand co-infection patterns in wildlife. By accounting for ecological, behavioural, and host-specific characteristics, it provides perspectives on the factors driving inter-individual variability of parasite community in a complex environment. Such an integrative approach advances our understanding of host-parasite systems and underscores the importance of considering both individual and environmental determinants to unravel co-infection mechanisms. The interplay between exposure and susceptibility factors, such as the influence of livestock presence, host activity, age-specific immune responses and between-parasite interactions, demonstrates the complexity of parasite-host relationships. Future research should expand on this framework by incorporating longitudinal studies to capture temporal dynamics, as well as exploring interactions at a finer scale, such as parasite-specific traits or immune pathways. Quantifying the relative contributions of these factors across different ecosystems could provide deeper insights into generalizable patterns of co-infection. Using recently developed hierarchical method, this work offers the groundwork for broader ecological investigations into parasite community dynamics, offering a foundation to better understand how environmental, behavioural, and physiological factors intersect in shaping parasite communities.

## Acknowledgements

This study was conducted as part of a PhD funded by Université Paul Sabatier and the doctoral school Sciences Ecologiques, Vétérinaire, Agronomiques & Bioingéniérie (SEVAB). We would thank everyone, technicians, researchers, veterinarians and volunteers that participated to the data record and the capture of animals. We also thank the PhyloPic community for their contribution to make available free naturalistic illustrations used in our graphical abstract.

## Author’s contribution

The conceptualization of the study has been done by CP, EGF, FB, HV and VB. Administration and funding support were managed by EGF and HV. Roe deer captures were performed by AB, FB, GLL, HV, JM, VB and YC. The livestock presence record was performed by BL, JM and YC. The data cleaning was realized by AB, FB and YC. ADN extractions were performed by SM and XB.

Parasite or antibody detections were performed by:

- JM and AB for gastrointestinal parasites
- FB, LM and CB for *Babesia* spp.
- FB, ACL, CR for *Bartonella* spp.
- VO and TB for *Borrelia* spp.
- CD and JI for Hepatitis E virus
- IV and DA for *Toxoplasma gondii*
- SM and XB for *Anaplasma phagocytophilum*
- LXN for *Mycoplasma* spp.

Methodology and statistical analysis were led by FB with the help of VS. FB wrote the original draft of the article with a main reviewing and edition by CP, EGF, HV and VB and a great help of VS and LM.

## Ethical standards

All protocols for capturing and marking roe deer were reviewed and approved by the French Ministry for Research and Higher Education and the ethics committee n°115 (project numbers Apafis#39320 and Apafis#7880). More detailed descriptions of these methods can be found in Morellet *et al*. 2009, Debeffe *et al*. 2013, and Bonnot *et al*. 2017, 2018. The laboratory Behaviour and Ecology of Wild Mammals ensures that the protocol complies with all animal welfare requirements and updates its protocol every year.

## References

Abbate, J. L, Galan, M, Razzauti, M, Sironen, T, Voutilainen, L, Henttonen, H, Gasqui, P, Cosson, J.-F. and Charbonnel, N (2024). Pathogen community composition and co-infection patterns in a wild community of rodents. Peer Community Journal 4, e14. doi: 10.24072/pcjournal.370.

Ahmed, N, French, T, Rausch, S, Kühl, A, Hemminger, K, Dunay, I. R, Steinfelder, S. and Hartmann, S (2017). Toxoplasma Co-infection Prevents Th2 Differentiation and Leads to a Helminth-Specific Th1 Response. Frontiers in Cellular and Infection Microbiology 7, 341. doi: 10.3389/fcimb.2017.00341.

Arneberg, P, Skorping, A, Grenfell, B. and Read, A. F (1998). Host densities as determinants of abundance in parasite communities. Proceedings of the Royal Society of London. Series B: Biological Sciences 265, 1283–1289. doi: 10.1098/rspb.1998.0431.

Aubert, D, Ajzenberg, D, Richomme, C, Gilot-Fromont, E, Terrier, M. E, de Gevigney, C, Game, Y, Maillard, D, Gibert, P, Dardé, M. L. and Villena, I (2010). Molecular and biological characteristics of Toxoplasma gondii isolates from wildlife in France. Veterinary Parasitology 171, 346–349. doi: 10.1016/j.vetpar.2010.03.033.

Barron, D. G, Gervasi, S. S, Pruitt, J. N. and Martin, L. B (2015). Behavioral competence: how host behaviors can interact to influence parasite transmission risk. Current opinion in behavioral sciences 6, 35–40. doi: 10.1016/j.cobeha.2015.08.002.

Barros, M, Cabezón, O, Dubey, J. P, Almería, S, Ribas, M. P, Escobar, L. E, Ramos, B. and Medina-Vogel, G (2018). Toxoplasma gondii infection in wild mustelids and cats across an urban-rural gradient. PLOS ONE 13, e0199085. doi: 10.1371/journal.pone.0199085.

Beaumelle, C, Redman, E, Verheyden, H, Jacquiet, P, Bégoc, N, Veyssière, F, Benabed, S, Cargnelutti, B, Lourtet, B, Poirel, M.-T, de Rijke, J, Yannic, G, Gilleard, J. S. and Bourgoin, G (2022). Generalist nematodes dominate the nemabiome of roe deer in sympatry with sheep at a regional level. International Journal for Parasitology 52, 751– 761. doi: 10.1016/j.ijpara.2022.07.005.

Benoit, L, Hewison, A. J. M, Coulon, A, Debeffe, L, Grémillet, D, Ducros, D, Cargnelutti, B, Chaval, Y. and Morellet, N (2020). Accelerating across the landscape: The energetic costs of natal dispersal in a large herbivore. Journal of Animal Ecology 89, 173–185. doi: 10.1111/1365-2656.13098.

Billeter, S. A, Levy, M. G, Chomel, B. B. and Breitschwerdt, E. B (2008). Vector transmission of Bartonella species with emphasis on the potential for tick transmission. Medical and Veterinary Entomology 22, 1–15. doi: 10.1111/j.1365-2915.2008.00713.x.

Body, G., Ferté, H., Gaillard, J.-M., Delorme, D., Klein, F. and Gilot-Fromont, E. (2011). Population density and phenotypic attributes influence the level of nematode parasitism in roe deer. Oecologia 167, 635–646.

Bohn, S. J, Webber, Q. M. R, Florko, K. R. N, Paslawski, K. R, Peterson, A. M, Piche, J. E, Menzies, A. K. and Willis, C. K. R (2017). Personality predicts ectoparasite abundance in an asocial sciurid. Ethology 123, 761–771. doi: 10.1111/eth.12651.

Bonnot, N, Verheyden, H, Blanchard, P, Cote, J, Debeffe, L, Cargnelutti, B, Klein, F, Hewison, A. J. M. and Morellet, N (2015). Interindividual variability in habitat use: evidence for a risk management syndrome in roe deer? Behavioral Ecology 26, 105–114. doi: 10.1093/beheco/aru169.

Bonnot, N. C, Goulard, M, Hewison, A. J. M, Cargnelutti, B, Lourtet, B, Chaval, Y. and Morellet, N (2018). Boldness-mediated habitat use tactics and reproductive success in a wild large herbivore. Animal Behaviour 145, 107–115. doi: 10.1016/j.anbehav.2018.09.013.

Bonte, D, Van Dyck, H, Bullock, J. M, Coulon, A, Delgado, M, Gibbs, M, Lehouck, V, Matthysen, E, Mustin, K, Saastamoinen, M, Schtickzelle, N, Stevens, V. M, Vandewoestijne, S, Baguette, M, Barton, K, Benton, T. G, Chaput-Bardy, A, Clobert, J, Dytham, C, Hovestadt, T, Meier, C. M, Palmer, S. C. F, Turlure, C. and Travis, J. M. J (2012). Costs of dispersal. Biological Reviews 87, 290–312. doi: 10.1111/j.1469-185X.2011.00201.x.

Brehm, A. M., Assis, V. R., Martin, L. B. and Orrock, J. L. Individual variation underlies large-scale patterns: Host conditions and behavior affect parasitism. Ecology n/a, e4478. doi: 10.1002/ecy.4478.

Bueno, C, Ruckstuhl, K. E, Arrigo, N, Aivaz, A. N. and Neuhaus, P (2012). Impacts of cattle grazing on small-rodent communities: an experimental case study. Canadian Journal of Zoology 90, 22–30. doi: 10.1139/z11-108.

Burt, W. H (1943). Territoriality and Home Range Concepts as Applied to Mammals. Journal of Mammalogy 24, 346–352. doi: 10.2307/1374834.

Carbillet, J., Hollain, M., Rey, B., Palme, R., Pellerin, M., Regis, C., Geffré, A., Duhayer, J., Pardonnet, S., Debias, F., Merlet, J., Lemaître, J.-F., Verheyden, H. and Gilot-Fromont, E. (2023). Age and spatio-temporal variations in food resources modulate stress-immunity relationships in three populations of wild roe deer. General and Comparative Endocrinology 330, 114141. doi: 10.1016/j.ygcen.2022.114141.

Cattadori, I. M, Boag, B. and Hudson, P. J (2008). Parasite co-infection and interaction as drivers of host heterogeneity. International Journal for Parasitology 38, 371–380. doi: 10.1016/j.ijpara.2007.08.004.

Chastagner, A, Pion, A, Verheyden, H, Lourtet, B, Cargnelutti, B, Picot, D, Poux, V, Bard, É, Plantard, O, McCoy, K. D, Leblond, A, Vourc’h, G. and Bailly, X (2017). Host specificity, pathogen exposure, and superinfections impact the distribution of Anaplasma phagocytophilum genotypes in ticks, roe deer, and livestock in a fragmented agricultural landscape. Infection, Genetics and Evolution 55, 31–44. doi: 10.1016/j.meegid.2017.08.010.

Cherry, N. A., Maggi, R. G., Cannedy, A. L. and Breitschwerdt, E. B. (2009). PCR detection of *Bartonella bovis* and *Bartonella henselae* in the blood of beef cattle. Veterinary Microbiology 135, 308–312. doi: 10.1016/j.vetmic.2008.09.063.

Cheynel, L., Lemaître, J.-F., Gaillard, J.-M., Rey, B., Bourgoin, G., Ferté, H., Jégo, M., Débias, F., Pellerin, M., Jacob, L. and Gilot-Fromont, E. (2017). Immunosenescence patterns differ between populations but not between sexes in a long-lived mammal. Scientific Reports 7, 13700. doi: 10.1038/s41598-017-13686-5.

Chirichella, R, Apollonio, M, Pokorny, B. and De Marinis, A. M (2023). Sex-specific impact of tooth wear on senescence in a low-dimorphic mammal species: The European roe deer (Capreolus capreolus). Journal of Zoology 319, 210–220. doi: 10.1111/jzo.13038.

Civitello, D. J. and Rohr, J. R (2014). Disentangling the effects of exposure and susceptibility on transmission of the zoonotic parasite *S chistosoma mansoni*. Journal of Animal Ecology 83, 1379–1386. doi: 10.1111/1365-2656.12222.

Cox, F. E. G (2001). Concomitant infections, parasites and immune responses. Parasitology 122, S23–S38. doi: 10.1017/S003118200001698X.

Cross, P. C, Drewe, J, Patrek, V, Pearce, G, Samuel, M. D. and Delahay, R. J (2009). Wildlife Population Structure and Parasite Transmission: Implications for Disease Management. In Management of Disease in Wild Mammals (ed. Delahay, R. J, Smith, G. C, and Hutchings, M. R.), pp. 9–29. Springer Japan, Tokyo doi: 10.1007/978-4-431-77134-0_2.

Dallas, T. A, Laine, A.-L. and Ovaskainen, O (2019). Detecting parasite associations within multi-species host and parasite communities. Proceedings of The Royal Society B: Biological Sciences 286,. doi: 10.1098/rspb.2019.1109.

de Cock, M. P, de Vries, A, Fonville, M, Esser, H. J, Mehl, C, Ulrich, R. G, Joeres, M, Hoffmann, D, Eisenberg, T, Schmidt, K, Hulst, M, van der Poel, W. H. M, Sprong, H. and Maas, M (2023). Increased rat-borne zoonotic disease hazard in greener urban areas. Science of The Total Environment 896, 165069. doi: 10.1016/j.scitotenv.2023.165069.

De La Peña, E, Martín, J, Barja, I, Pérez-Caballero, R, Acosta, I. and Carranza, J (2020). Immune challenge of mating effort: steroid hormone profile, dark ventral patch and parasite burden in relation to intrasexual competition in male Iberian red deer. Integrative Zoology 15, 262–275. doi: 10.1111/1749-4877.12427.

Debeffe, L, Morellet, N, Cargnelutti, B, Lourtet, B, Bon, R, Gaillard, J.-M. and Mark Hewison, A. J (2012). Condition-dependent natal dispersal in a large herbivore: heavier animals show a greater propensity to disperse and travel further. Journal of Animal Ecology 81, 1327–1327. doi: 10.1111/j.1365-2656.2012.02014.x.

Delsart, M, Fablet, C, Rose, N, Répérant, J.-M, Blaga, R, Dufour, B. and Pol, F (2022). Descriptive Epidemiology of the Main Internal Parasites on Alternative Pig Farms in France. Journal of Parasitology 108, 306–321. doi: 10.1645/21-126.

Destoumieux-Garzón, D, Mavingui, P, Boetsch, G, Boissier, J, Darriet, F, Duboz, P, Fritsch, C, Giraudoux, P, Le Roux, F, Morand, S, Paillard, C, Pontier, D, Sueur, C. and Voituron, Y (2018). The One Health Concept: 10 Years Old and a Long Road Ahead. Frontiers in Veterinary Science 5, 14. doi: 10.3389/fvets.2018.00014.

Djokic, V., Blaga, R., Aubert, D., Durand, B., Perret, C., Geers, R., Ducry, T., Vallee, I., Djakovic, O. D., Mzabi, A., Villena, I. and Boireau, P. (2016). Toxoplasma gondii infection in pork produced in France. Parasitology 143, 557–567. doi: 10.1017/S0031182015001870.

Dougherty, E. R, Seidel, D. P, Carlson, C. J, Spiegel, O. and Getz, W. M (2018). Going through the motions: incorporating movement analyses into disease research. Ecology Letters 21, 588–604. doi: 10.1111/ele.12917.

Drazenovich, N, Foley, J. and Brown, R. N (2006). Use of Real-Time Quantitative PCR Targeting the msp2 Protein Gene to Identify Cryptic Anaplasma phagocytophilum Infections in Wildlife and Domestic Animals. Vector-Borne and Zoonotic Diseases 6, 83–90. doi: 10.1089/vbz.2006.6.83.

Dunn, J. C, Cole, E. F. and Quinn, J. L (2011). Personality and parasites: sex-dependent associations between avian malaria infection and multiple behavioural traits. Behavioral Ecology and Sociobiology 65, 1459–1471. doi: 10.1007/s00265-011-1156-8.

Ezenwa, V. O (2016). Helminth–microparasite co-infection in wildlife: lessons from ruminants, rodents and rabbits. Parasite Immunology 38, 527–534. doi: 10.1111/pim.12348.

Ezenwa, V. O. and Jolles, A. E (2015). Opposite effects of anthelmintic treatment on microbial infection at individual versus population scales. Science 347, 175–177. doi: 10.1126/science.1261714.

Ezenwa, V. O, Etienne, R. S, Luikart, G, Beja-Pereira, A. and Jolles, A. E (2010). Hidden Consequences of Living in a Wormy World: Nematode-Induced Immune Suppression Facilitates Tuberculosis Invasion in African Buffalo. The American Naturalist 176, 613– 624. doi: 10.1086/656496.

Fernández-i-Marín, X (2016). ggmcmc: Analysis of MCMC Samples and Bayesian Inference. Journal of Statistical Software 70, 1–20. doi: 10.18637/jss.v070.i09.

Ferrari, N., Cattadori, I. M., Nespereira, J., Rizzoli, A. and Hudson, P. J. (2004). The role of host sex in parasite dynamics: field experiments on the yellow-necked mouse Apodemus flavicollis. Ecology Letters 7, 88–94. doi: 10.1046/j.1461-0248.2003.00552.x.

Foo, Y. Z, Nakagawa, S, Rhodes, G. and Simmons, L. W (2017). The effects of sex hormones on immune function: a meta-analysis. Biological Reviews 92, 551–571. doi: 10.1111/brv.12243.

Fox, N. J, Marion, G, Davidson, R. S, White, P. C. L. and Hutchings, M. R (2013). Modelling Parasite Transmission in a Grazing System: The Importance of Host Behaviour and Immunity. PLoS ONE 8, e77996. doi: 10.1371/journal.pone.0077996.

Gardner, E. M. and Murasko, D. M (2002). Age-related changes in Type 1 and Type 2 cytokine production in humans. Biogerontology 3, 271–290. doi: 10.1023/A:1020151401826.

Gasparoni, A, Ciardelli, L, Avanzini, A, Castellazzi, A. M, Carini, R, Rondini, G. and Chirico, G (2003). Age-Related Changes in Intracellular Th1/Th2 Cytokine Production, Immunoproliferative T Lymphocyte Response and Natural Killer Cell Activity in Newborns, Children and Adults. Neonatology 84, 297–303. doi: 10.1159/000073638.

Gervais, L, Hewison, A. J. M, Morellet, N, Bernard, M, Merlet, J, Cargnelutti, B, Chaval, Y, Pujol, B. and Quéméré, E (2020). Pedigree-free quantitative genetic approach provides evidence for heritability of movement tactics in wild roe deer. Journal of Evolutionary Biology 33, 595–607. doi: 10.1111/jeb.13594.

González, D. A, Díaz, B. B, Rodríguez Pérez, M. del C, Hernández, A. G, Chico, B. N. D. and de León, A. C (2010). Sex hormones and autoimmunity. Immunology Letters 133, 6–13. doi: 10.1016/j.imlet.2010.07.001.

Goossens, S., Wybouw, N., Van Leeuwen, T. and Bonte, D. (2020). The physiology of movement. Movement Ecology 8, 5. doi: 10.1186/s40462-020-0192-2.

Gortázar, C, Ruiz-Fons, J. F. and Höfle, U (2016). Infections shared with wildlife: an updated perspective. European Journal of Wildlife Research 62, 511–525. doi: 10.1007/s10344-016-1033-x.

Gotteland, C, Chaval, Y, Villena, I, Galan, M, Geers, R, Aubert, D, Poulle, M.-L, Charbonnel, N. and Gilot-Fromont, E (2014). Species or local environment, what determines the infection of rodents by *Toxoplasma gondii*? Parasitology 141, 259–268. doi: 10.1017/S0031182013001522.

Gracia, M. J., Calvete, C., Estrada, R., Castillo, J. A., Peribáñez, M. A. and Lucientes, J. (2013). Survey of flea infestation in cats in Spain. Medical and Veterinary Entomology 27, 175–180. doi: 10.1111/j.1365-2915.2012.01027.x.

Graham, A. L (2008). Ecological rules governing helminth–microparasite coinfection. Proceedings of the National Academy of Sciences 105, 566–570. doi: 10.1073/pnas.0707221105.

Gubler, D. J, Reiter, P, Ebi, K. L, Yap, W, Nasci, R. and Patz, J. A (2001). Climate variability and change in the United States: potential impacts on vector- and rodent-borne diseases. Environmental Health Perspectives 109, 223–233. doi: 10.1289/ehp.109-1240669.

Hawley, D. M, Etienne, R. S, Ezenwa, V. O. and Jolles, A. E (2011). Does animal behavior underlie covariation between hosts’ exposure to infectious agents and susceptibility to infection? Implications for disease dynamics. Integrative and Comparative Biology 51, 528–539. doi: 10.1093/icb/icr062.

Hewison, A. J. M, Vincent, J. P. and Reby, D (1998). Social organisation of European roe deer. In The European roe deer : The biology of success, p. Scandinavian University Press.

Hoffmann, S., Horak, I. G., Bennett, N. C. and Lutermann, H. (2016). Evidence for interspecific interactions in the ectoparasite infracommunity of a wild mammal. Parasites & Vectors 9, 58. doi: 10.1186/s13071-016-1342-7.

Hofmann-Lehmann, R, Meli, M. L, Dreher, U. M, Gönczi, E, Deplazes, P, Braun, U, Engels, M, Schüpbach, J, Jörger, K, Thoma, R, Griot, C, Stärk, K. D. C, Willi, B, Schmidt, J, Kocan, K. M. and Lutz, H (2004). Concurrent Infections with Vector-Borne Pathogens Associated with Fatal Hemolytic Anemia in a Cattle Herd in Switzerland. Journal of Clinical Microbiology 42, 3775–3780. doi: 10.1128/JCM.42.8.3775-3780.2004.

Horcajada-Sánchez, F, Navarro-Castilla, Á, Boadella, M. and Barja, I (2018). Influence of livestock, habitat type, and density of roe deer (Capreolus capreolus) on parasitic larvae abundance and infection seroprevalence in wild populations of roe deer from central Iberian Peninsula. Mammal Research 63, 213–222. doi: 10.1007/s13364-018-0354-4.

Jensen, W. A, Lappin, M. R, Kamkar, S and Reagan, W. J (2001). Use of a polymerase chain reaction assay to detect and differentiate two strains of *Haemobartonella felis* in naturally infected cats. American Journal of Veterinary Research 62, 604–608. doi: 10.2460/ajvr.2001.62.604.

Johnson, P. T. J, Wood, C. L, Joseph, M. B, Preston, D. L, Haas, S. E. and Springer, Y. P (2016). Habitat heterogeneity drives the host-diversity-begets-parasite-diversity relationship: evidence from experimental and field studies. Ecology Letters 19, 752–761. doi: 10.1111/ele.12609.

Jones, K. E, Patel, N. G, Levy, M. A, Storeygard, A, Balk, D, Gittleman, J. L. and Daszak, P (2008). Global trends in emerging infectious diseases. Nature 451, 990–993. doi: 10.1038/nature06536.

Kahl, O. and Gray, J. S (2023). The biology of Ixodes ricinus with emphasis on its ecology. Ticks and Tick-borne Diseases 14, 102114. doi: 10.1016/j.ttbdis.2022.102114.

Keegan, S. P, Pedersen, A. B. and Fenton, A (2024). The impact of within-host coinfection interactions on between-host parasite transmission dynamics varies with spatial scale. Proceedings of the Royal Society B: Biological Sciences 291, 20240103. doi: 10.1098/rspb.2024.0103.

Kiffner, C, Lödige, C, Alings, M, Vor, T. and Rühe, F (2011). Body-mass or sex-biased tick parasitism in roe deer (Capreolus capreolus)? A GAMLSS approach. Medical and Veterinary Entomology 25, 39–45. doi: 10.1111/j.1365-2915.2010.00929.x.

Klein, S. L (2000). The effects of hormones on sex differences in infection: from genes to behavior. Neuroscience & Biobehavioral Reviews 24, 627–638. doi: 10.1016/S0149-7634(00)00027-0.

Knowles, S. C. L., Fenton, A., Petchey, O. L., Jones, T. R., Barber, R. and Pedersen, A. B. (2013). Stability of within-host–parasite communities in a wild mammal system. Proceedings of the Royal Society B: Biological Sciences 280, 20130598. doi: 10.1098/rspb.2013.0598.

Kovacs, E. J, Duffner, L. A. and Plackett, T. P (2004). Immunosuppression after injury in aged mice is associated with a TH1–TH2 shift, which can be restored by estrogen treatment. Mechanisms of Ageing and Development 125, 121–123. doi: 10.1016/j.mad.2003.11.007.

Krasnov, B. R., Stanko, M. and Morand, S. (2010). Competition, facilitation or mediation via host? Patterns of infestation of small European mammals by two taxa of haematophagous arthropods. Ecological Entomology 35, 37–44. doi: 10.1111/j.1365-2311.2009.01153.x.

Lehmann, T, Graham, D. H, Dahl, E, Sreekumar, C, Launer, F, Corn, J. L, Gamble, H. R. and Dubey, J. P (2003). Transmission dynamics of *Toxoplasma gondii* on a pig farm. *Infection*, Genetics and Evolution 3, 135–141. doi: 10.1016/S1567-1348(03)00067-4.

Linnell, J. D. C., Cretois, B., Nilsen, E. B., Rolandsen, C. M., Solberg, E. J., Veiberg, V., Kaczensky, P., Van Moorter, B., Panzacchi, M., Rauset, G. R. and Kaltenborn, B. (2020). The challenges and opportunities of coexisting with wild ungulates in the human-dominated landscapes of Europe’s Anthropocene. Biological Conservation 244, 108500. doi: 10.1016/j.biocon.2020.108500.

Lutermann, H., Fagir, D. M. and Bennett, N. C. (2015). Complex interactions within the ectoparasite community of the eastern rock sengi (*Elephantulus myurus*). International Journal for Parasitology: Parasites and Wildlife 4, 148–158. doi: 10.1016/j.ijppaw.2015.02.001.

Maizels, R. M, Balic, A, Gomez-Escobar, N, Nair, M, Taylor, M. D. and Allen, J. E (2004). Helminth parasites – masters of regulation. Immunological Reviews 201, 89–116. doi: 10.1111/j.0105-2896.2004.00191.x.

Malagnino, A, Marchand, P, Garel, M, Cargnelutti, B, Itty, C, Chaval, Y, Hewison, A. J. M, Loison, A. and Morellet, N (2021). Do reproductive constraints or experience drive age-dependent space use in two large herbivores? Animal Behaviour 172, 121–133. doi: 10.1016/j.anbehav.2020.12.004.

McArdle, A. J, Turkova, A. and Cunnington, A. J (2018). When do co-infections matter? Current Opinion in Infectious Diseases 31, 209–215. doi: 10.1097/QCO.0000000000000447.

McCauley, D. J, Keesing, F, Young, T. and Dittmar, K (2008). Effects of the removal of large herbivores on fleas of small mammals. Journal of Vector Ecology 33, 263–268. doi: 10.3376/1081-1710-33.2.263.

Moutailler, S, Valiente Moro, C, Vaumourin, E, Michelet, L, Tran, F. H, Devillers, E, Cosson, J.-F, Gasqui, P, Van, V. T, Mavingui, P, Vourc’h, G. and Vayssier-Taussat, M (2016). Co-infection of Ticks: The Rule Rather Than the Exception. PLOS Neglected Tropical Diseases 10, e0004539. doi: 10.1371/journal.pntd.0004539.

Nouvel, L. X, Hygonenq, M.-C, Catays, G, Martinelli, E, Le Page, P, Collin, É, Inokuma, H, Schelcher, F, Citti, C. and Maillard, R (2019). First detection of Mycoplasma wenyonii in France: Identification, evaluation of the clinical impact and development of a new specific detection assay. *Comparative Immunology*, Microbiology and Infectious Diseases 63, 148–153. doi: 10.1016/j.cimid.2019.01.010.

Obiegala, A, Król, N, Oltersdorf, C, Nader, J. and Pfeffer, M (2017). The enzootic life-cycle of Borrelia burgdorferi (sensu lato) and tick-borne rickettsiae: an epidemiological study on wild-living small mammals and their ticks from Saxony, Germany. Parasites & Vectors 10, 115. doi: 10.1186/s13071-017-2053-4.

Ollivier, V, Choquet, R, Gamble, A, Bastien, M, Combes, B, Gilot-Fromont, E, Pellerin, M, Gaillard, J.-M, Lemaître, J.-F, Verheyden, H. and Boulinier, T (2023). Temporal dynamics of antibody level against Lyme disease bacteria in roe deer: Tale of a sentinel? Ecology and Evolution 13, e10414. doi: 10.1002/ece3.10414.

Ovaskainen, O. and Abrego, N (2020). Joint Species Distribution Modelling: With Applications in R. Cambridge University Press, Cambridge doi: 10.1017/9781108591720.

Ovaskainen, O, Tikhonov, G, Norberg, A, Guillaume Blanchet, F, Duan, L, Dunson, D, Roslin, T. and Abrego, N (2017). How to make more out of community data? A conceptual framework and its implementation as models and software. Ecology Letters 20, 561–576. doi: 10.1111/ele.12757.

Parker, I. M, Saunders, M, Bontrager, M, Weitz, A. P, Hendricks, R, Magarey, R, Suiter, K. and Gilbert, G. S (2015). Phylogenetic structure and host abundance drive disease pressure in communities. Nature 520, 542–544. doi: 10.1038/nature14372.

Pato, F. J, Vázquez, L, Díez-Baños, N, López, C, Sánchez-Andrade, R, Fernández, G, Díez-Baños, P, Panadero, R, Díaz, P. and Morrondo, P (2013). Gastrointestinal nematode infections in roe deer (Capreolus capreolus) from the NW of the Iberian Peninsula: Assessment of some risk factors. Veterinary Parasitology 196, 136–142. doi: 10.1016/j.vetpar.2013.01.027.

Patterson, L. D. and Schulte-Hostedde, A. I (2011). Behavioural correlates of parasitism and reproductive success in male eastern chipmunks, Tamias striatus. Animal Behaviour 81, 1129–1137. doi: 10.1016/j.anbehav.2011.02.016.

Pearce, J. and Ferrier, S (2000). Evaluating the predictive performance of habitat models developed using logistic regression. Ecological Modelling 133, 225–245. doi: 10.1016/S0304-3800(00)00322-7.

Pedersen, A. B. and Antonovics, J. (2013). Anthelmintic treatment alters the parasite community in a wild mouse host. Biology Letters 9, 20130205. doi: 10.1098/rsbl.2013.0205.

Perkins, S. E., Cattadori, I. M., Tagliapietra, V., Rizzoli, A. P. and Hudson, P. J. (2003). Empirical evidence for key hosts in persistence of a tick-borne disease. International Journal for Parasitology 33, 909–917. doi: 10.1016/S0020-7519(03)00128-0.

Poulin, R (2013). Parasite manipulation of host personality and behavioural syndromes. Journal of Experimental Biology 216, 18–26. doi: 10.1242/jeb.073353.

Qasem, L, Cardew, A, Wilson, A, Griffiths, I, Halsey, L. G, Shepard, E. L. C, Gleiss, A. C. and Wilson, R (2012). Tri-Axial Dynamic Acceleration as a Proxy for Animal Energy Expenditure; Should We Be Summing Values or Calculating the Vector? PLoS ONE 7, e31187. doi: 10.1371/journal.pone.0031187.

Rampersad, J. N, Watkins, J. D, Samlal, M. S, Deonanan, R, Ramsubeik, S. and Ammons, D. R (2005). A nested-PCR with an Internal Amplification Control for the detection and differentiation of Bartonella henselae and B. clarridgeiae: An examination of cats in Trinidad. BMC Infectious Diseases 5, 63. doi: 10.1186/1471-2334-5-63.

Raynaud, J.-P, William, G. and Brunault, G (1970). Etude de l’efficacité d’une technique de coproscopie quantitative pour le diagnostic de routine et le contrôle des infestations parasitaires des bovins, ovins, équins et porcins. Annales de Parasitologie Humaine et Comparée 45, 321–342. doi: 10.1051/parasite/1970453321.

Rohr, J. R, Civitello, D. J, Halliday, F. W, Hudson, P. J, Lafferty, K. D, Wood, C. L. and Mordecai, E. A (2019). Towards common ground in the biodiversity–disease debate. Nature Ecology & Evolution 4, 24–33. doi: 10.1038/s41559-019-1060-6.

Rolff, J. (2002). Bateman’s principle and immunity. Proceedings of the Royal Society of London. Series B: Biological Sciences 269, 867–872. doi: 10.1098/rspb.2002.1959.

Santicchia, F, Romeo, C, Ferrari, N, Matthysen, E, Vanlauwe, L, Wauters, L. A. and Martinoli, A (2019). The price of being bold? Relationship between personality and endoparasitic infection in a tree squirrel. Mammalian Biology 97, 1–8. doi: 10.1016/j.mambio.2019.04.007.

Sevila, J, Richomme, C, Hoste, H, Candela, M. G, Gilot-Fromont, E, Rodolakis, A, Cebe, N, Picot, D, Merlet, J. and Verheyden, H (2014). Does land use within the home range drive the exposure of roe deer (Capreolus capreolus) to two abortive pathogens in a rural agro-ecosystem? Acta Theriologica 59, 571–581. doi: 10.1007/s13364-014-0197-6.

Sih, A., Bell, A. and Johnson, J. C. (2004). Behavioral syndromes: an ecological and evolutionary overview. Trends in Ecology & Evolution 19, 372–378. doi: 10.1016/j.tree.2004.04.009.

Sih, A., Spiegel, O., Godfrey, S., Leu, S. and Bull, C. M. (2018). Integrating social networks, animal personalities, movement ecology and parasites: a framework with examples from a lizard. Animal Behaviour 136, 195–205. doi: 10.1016/j.anbehav.2017.09.008.

Sisson, D, Beechler, B, Jabbar, A, Jolles, A. and Hufschmid, J (2023). Epidemiology of *Anaplasma marginale* and *Anaplasma centrale* infections in African buffalo (*Syncerus caffer*) from Kruger National Park, South Africa. International Journal for Parasitology: Parasites and Wildlife 21, 47–54. doi: 10.1016/j.ijppaw.2023.04.005.

Skotarczak, B. and Adamska, M. (2005). Detection of Bartonella DNA in roe deer (Capreolus capreolus) and in ticks removed from deer. European Journal of Wildlife Research 51, 287–290. doi: 10.1007/s10344-005-0112-1.

Smith, K. F, Goldberg, M, Rosenthal, S, Carlson, L, Chen, J, Chen, C. and Ramachandran, S (2014). Global rise in human infectious disease outbreaks. Journal of The Royal Society Interface 11, 20140950. doi: 10.1098/rsif.2014.0950.

Storey, G. W. and Phillips, R. A (1985). The survival of parasite eggs throughout the soil profile. Parasitology 91, 585–590. doi: 10.1017/S003118200006282X.

Sweeny, A. R. and Albery, G. F (2022). Exposure and susceptibility: The Twin Pillars of infection. Functional Ecology 36, 1713–1726. doi: 10.1111/1365-2435.14065.

Teitelbaum, C. S, Huang, S, Hall, R. J. and Altizer, S (2018). Migratory behaviour predicts greater parasite diversity in ungulates. Proceedings of the Royal Society B: Biological Sciences 285, 20180089. doi: 10.1098/rspb.2018.0089.

Telfer, S, Lambin, X, Birtles, R. J, Beldomenico, P. M, Burthe, S. J, Paterson, S. and Begon, M (2010). Species Interactions in a Parasite Community Drive Infection Risk in a Wildlife Population. Science 330, 243–246. doi: 10.1126/science.1190333.

Tikhonov, G, Opedal, Ø. H, Abrego, N, Lehikoinen, A, de Jonge, M. M. J, Oksanen, J. and Ovaskainen, O (2020). Joint species distribution modelling with the r -package H msc. Methods in Ecology and Evolution 11, 442–447. doi: 10.1111/2041-210X.13345.

Tjur, T (2009). Coefficients of Determination in Logistic Regression Models—A New Proposal: The Coefficient of Discrimination. The American Statistician 63, 366–372. doi: 10.1198/tast.2009.08210.

Tricou, V., Bouscaillou, J., Laghoe-Nguembe, G.-L., Béré, A., Konamna, X., Sélékon, B., Nakouné, E., Kazanji, M. and Komas, N. P. (2020). Hepatitis E virus outbreak associated with rainfall in the Central African Republic in 2008-2009. BMC Infectious Diseases 20, 260. doi: 10.1186/s12879-020-04961-4.

Turner, W. C. and Getz, W. M (2010). SEASONAL AND DEMOGRAPHIC FACTORS INFLUENCING GASTROINTESTINAL PARASITISM IN UNGULATES OF ETOSHA NATIONAL PARK. Journal of Wildlife Diseases 46, 1108–1119. doi: 10.7589/0090-3558-46.4.1108.

Vanden Broecke, B, Bernaerts, L, Ribas, A, Sluydts, V, Mnyone, L, Matthysen, E. and Leirs, H (2021). Linking behavior, Co-infection Patterns, and Viral Infection Risk With the Whole Gastrointestinal Helminth Community Structure in Mastomys natalensis. Frontiers in Veterinary Science 8, 669058. doi: 10.3389/fvets.2021.669058.

Vanden Broecke, B, Tafompa, P. J. J, Mwamundela, B. E, Bernaerts, L, Ribas, A, Mnyone, L. L, Leirs, H. and Mariën, J (2023). Drivers behind co-occurrence patterns between pathogenic bacteria, protozoa, and helminths in populations of the multimammate mouse, Mastomys natalensis. Acta Tropica 243, 106939. doi: 10.1016/j.actatropica.2023.106939.

Verheyden, H, Richomme, C, Sevila, J, Merlet, J, Lourtet, B, Chaval, Y. and Hoste, H (2020). Relationship between the excretion of eggs of parasitic helminths in roe deer and local livestock density. Journal of Helminthology 94, e159. doi: 10.1017/S0022149X20000449.

Viney, M. E. and Graham, A. L (2013). Patterns and Processes in Parasite Co-Infection. In Advances in Parasitology, pp. 321–369. Elsevier doi: 10.1016/B978-0-12-407706-5.00005-8.

Vor, T, Kiffner, C, Hagedorn, P, Niedrig, M. and Rühe, F (2010). Tick burden on European roe deer (Capreolus capreolus). Experimental and Applied Acarology 51, 405–417. doi: 10.1007/s10493-010-9337-0.

Wang, Y, Zhang, H, Li, Z, Gu, W, Lan, H, Hao, W, Ling, R, Li, H. and Harrison, T. J (2001). Detection of Sporadic Cases of Hepatitis E Virus (HEV) Infection in China Using Immunoassays Based on Recombinant Open Reading Frame 2 and 3 Polypeptides from HEV Genotype 4. Journal of Clinical Microbiology 39, 4370. doi: 10.1128/JCM.39.12.4370-4379.2001.

Waruiru, R. M., Kyvsgaard, N. C., Thamsborg, S. M., Nansen, P., Bøgh, H. O., Munyua, W. K. and Gathuma, J. M (2000). The Prevalence and Intensity of Helminth and Coccidial Infections in Dairy Cattle in Central Kenya. Veterinary Research Communications 24, 39–53. doi: 10.1023/A:1006325405239.

Watanabe, S (2010). Equations of states in singular statistical estimation. Neural Networks 23, 20–34. doi: 10.1016/j.neunet.2009.08.002.

Wilson, R. P, White, C. R, Quintana, F, Halsey, L. G, Liebsch, N, Martin, G. R. and Butler, P. J (2006). Moving towards acceleration for estimates of activity-specific metabolic rate in free-living animals: the case of the cormorant. Journal of Animal Ecology 75, 1081–1090. doi: 10.1111/j.1365-2656.2006.01127.x.

Worton, B. J (1989). Kernel Methods for Estimating the Utilization Distribution in Home-Range Studies. Ecology 70, 164–168. doi: 10.2307/1938423.

Zhioua, E, Aeschlimann, A. and Gern, L (1994). Infection of Field-Collected Ixodes ricinus (Acari: Ixodidae) Larvae with Borrelia burgdorferi in Switzerland. Journal of Medical Entomology 31, 763–766. doi: 10.1093/jmedent/31.5.763.

