## Supplementary Material for "Effects of host sex, age and behaviour on co-infection patterns in a wild ungulate"

### *Protocols for parasite and antibody detection*

#### *Anaplasma phagocytophilum* detection

The presence of *Anaplasma phagocytophilum* was detected by real-time quantitative PCR targeting a 122-bp fragment of the *msp2* gene using primers 903f (5'-AGTTTGACTGGAACACACCTGATC-3') and 1024r (5'-CTCGTAACCAATCTCAAGCTCAAC-3') and the dual labelled probe (939p 5'-FAM-TTAAGGACAACATGCTTGTAGCTATGGAAGGCA-BHQ1-3') (Drazenovich, Foley et Brown, 2006). Taqman qPCRs were performed in 96 well plates using a Bio-Rad CFX96 Real-time PCR thermocycler. Each reaction had a final volume of 20 µL. It included 1× SsoAdvanced™ universal Probes Supermix (Bio-Rad, Hercules, USA), 250 nM of probe, 1 µM of each primer (Eurofins Genomics, Ebersberg, Germany). We added 5 µL of genomic DNA extract from each whole blood sample and molecular biology grade water to reach the final reaction volume. Five µL of the negative extraction control of the genomic DNA series analysed and a negative control of qPCR reaction mixtures, containing 5 µL of molecular biology grade water in place of the DNA extract were included on each PCR plate. qPCR cycling conditions included a 3 min at 95 ◦C initial step for polymerase activation and DNA denaturation, followed by 40 three-step cycles of i) denaturation at 95 ◦C for 15 s and ii) annealing-extension at 60 ◦C for 30 s and iii) fluorescence measurement. The quantification cycle (Cq) was determined using Bio-Rad CFX Maestro software in Regression mode.

### *Borrelia* spp. antibodies detection

For GA samples, blotting papers were eluted in a DILBUF buffer overnight by placing a 1 $cm^2$ paper piece in a 2 mL tube with the buffer. Antibody levels against Bbsl in serum and blotting paper samples from roe deer were quantified using a modified commercial anti-Bbsl ELISA kit (Borrelia + VlsE IgG ELISA Kit; IBL International). The peroxidase-conjugated anti-human IgG antibodies in the kit were replaced with anti-deer IgG antibodies peroxidase-labelled antibodies to deer IgG (H +L); 04-31-06, KPL). Antibody levels were measured as optical density (OD) at 450 nm, and the serological status was determined from the OD distribution using mixture models in R software.

To identify the best model for OD distribution (seronegative vs. seropositive or three distributions due to individual differences), the Akaike information criterion (AIC) was used. The threshold for seropositivity was set based on the 95% confidence interval of the seronegative samples. This threshold was validated by immunoblots on 28 serum samples with different OD values to confirm antibody reactions to Bbsl markers (e.g., VlsE).

Finally, a correlation analysis between annual OD measurements from blood and blotting papers from the same individuals showed a strong correlation ($r^2 = 0.92$, $n = 20$), indicating the assay's stability.

### *Babesia venatorum* and *Babesia capreoli* detection

The presence of piroplasms *B. venatorum* and *B. capreoli* was detected by duplex real-time quantitative PCR. Primers common to both species (18Sq_fw 5'-GCAGTTAAAAAGCTCGTAGTTG-3' and 18SqEU1cap_rev 5'-CAAAAGTCTGCTTGAAACACTC-3') and double-dye probes (18S-Bvena_probe 5'-6-FAM-CTGCGTTATCGAGTTATTGACTCTT-BHQ1-3' and 18S-Bcapre_probe 5'-6-HEX-CGTGGTGTTAATATTGACTGATGTC-BHQ1-3') were specifically designed for this study to amplify and detect a 128 bp fragment within the V4 hypervariable region of 18S rRNA gene. The probes were labelled at the 5’ end with the reporter dye 6-Carboxyfluorescein (FAM) or 5-Hexachloro-fluorescein (HEX) for *B. venatorum* or *B. capreoli*, respectively. For both, the 3’ end was labelled with Black Hole Quencher-1 (BHQ1) (Eurogentec, Belgium).

Reactions were carried out in 10 µL reaction mixture containing 450 µM of each primer, 300 µM of each probe, 1X SsoAdvanced universal probes supermix (Bio-Rad, CA, USA) and 2 µL of DNA sample. A positive control consisting of DNA from *B. venatorum* C201 isolate mixed with DNA from *B. capreoli* 2770F6 isolate, and a negative control with water were added to each run. All qPCR assays were performed in the CFX Opus 96 Real-Time PCR instrument (Bio-Rad, CA, USA) with cycling conditions as follows: 95°C for 3 min, then 40 cycles of 15 s at 95°C and 30 s at 60°C. The quantification cycle (Cq) was determined using Bio-Rad CFX Maestro software in Regression mode.

### *Bartonella* spp. detection

The presence of *Bartonella* spp. DNA in samples was assessed by a nested PCR targeting the *16S-23S ribosomal RNA* intergenic spacer according to Rampersad *et al.* 2005. Primers P-bhenfa (5’-TCTTCGTTTCTCTTTCTTCA-3’) and P-benr1 (5’CAAGCGCGCGCTCTAACC-3’) were used for the external PCR and primers N-bhenf1A (5’-GATGATCCCAAGCCTTCTGGC-3’) and N-bhenr (5’-AACCAACTGAGCTACAAGCC-3’) for the internal PCR. PCR was performed in a final volume of 25 µL using 1X *Ex Taq* Buffer, 200 µM dNTP, 1 U *TaKaRa Ex Taq* (TaKaRa, Shiga, Japan), 0.8 µM of each primer and sterile water. Five microliters of DNA or amplified PCR products were added for external and internal PCR, respectively. *Bartonella henselae* DNAand sterile water were used as positive and negative controls, respectively. Thermal cycling conditions included an initial activation of Taq polymerase at 95°C for 30 s, followed by 39 cycles of a-15 s denaturation step at 94°C, a-30 s annealing step at 53°C (for external PCR) or 57°C (for internal PCR), and a-30 s elongation step at 72°C, followed by a final extension step at 72°C for 5 min. PCR amplification products were separated on a 1.8% agarose gel stained with ethidium bromide and visualised under ultraviolet light.

### *Mycoplasma* spp. detection

The presence of *Mycoplasma* spp. was assessed by PCR detection of DNA in samples according to the protocol used by Nouvel *et al.* 2019 targeting a portion of the 16S ribosomal RNA gene that is conserved in hemotropic mycoplasmas species, also named hemoplasmas (Jensen *et al.et al.* 2001) adapted for real time PCR (qPCR). Briefly, DNA was extracted from 200 µL of EDTA-blood samples using a QIAamp DNA Blood Mini Kit (QIAGEN), following manufacturer's instructions, eluted with 100 µl of buffer AE and stored at −20°C until use. RT-Quantitative PCR assays were performed on a LightCycler 96 thermocycler (Roche, Basel, Switzerland) with the following volumes for each sample analysed:  total volume of 20 µL, with 2 µL of DNA sample, 0.4 µL of each 10 µM primer (final concentration of 0.2 µM), 10 µL of SYBR Green I Master, 7.2 µL of PCR-grade water. Thermocycling programs were set on the same following conditions: pre-incubation step of 10 min at 95°C, 45 amplification cycles of 10 s at 95°C, 15s at 60°C and 15 s at 72°C with single acquisition, and a melting analysis phase of 10 s at 95°C, 1 min at 65°C and then 1 s heating at 97°C with continuous acquisition (5 readings per degree Celsius). Negative and positive PCR controls were systematically run along with samples and consisted respectively of 2 µL of water and 2 µL of standard dilutions of plasmid construct including the specific PCR target sequence.

### Coccidia, Strongylidae and *Nematodirus* spp. detection

The parasitological coprology protocol involves processing faecal samples to quantify parasite eggs using a McMaster counting slide. A hypersaline solution (200 g of salt in 600 ml of water) is prepared, with 42 ml used per 3 g of faeces, adjusting the solution if less faecal material is available. The faeces are first finely crushed with a mortar and pestle before gradually mixing with the saline solution to homogenise the sample. The mixture is filtered through a fine sieve (0.5 mm mesh), and the filtrate is further homogenised using a pipette.

The McMaster slide chambers are then filled with the prepared solution, ensuring no air bubbles are present. After allowing the slide to rest for at least two minutes to let parasite eggs float to the surface, the sample is examined under a microscope at 100x magnification, focusing on one column at a time. Eggs within the grid are counted and multiplied by 50 to estimate egg concentration and identify as Coccidia, Strongylidae, *Nematodirus* spp. or Trichuris. If discrepancies between grids occur, a new slide is prepared. When no eggs are found within the grids, the area outside the grid is examined, and the count is multiplied by 15.

### Hepatitis E Virus (HEV) detection

Fifty-five plasmas and 364 roe deer sera were collected from a population monitored between 2016 and 2023. Some animals were collected several times between different years. Blood samples were stored at -20°C prior to analysis. Four hundred and seven samples were analysed. Samples were extracted using the Altostar® technology and stored at -80°C before and after analysis. Indirect ELISA was then performed on 96 well plates in order to detect HEV antibodies in a qualitative way (IgM, IgG). 3 wells were used as negative control and 2 as positive control for every tested plate. Antibody capture was done in a solid phase after successive washings. The fluorescence was emitted at 450 nm by horseradish peroxidase and revealed by a chromogenic solution of Tetramethylbenzidine, enabling positive and negative wells to be differentiated. The colour obtained was proportional to the amount of complex captured.

*Toxoplasma gondii* detection

Blood serum samples were screened for *T. gondii* antibodies using the modified agglutination test (MAT), following the method outlined by Dubey and Desmonts (1987). Serum samples were initially diluted two-fold at a 1:6 starting dilution. Consistent with prior studies, samples with an agglutination titer of 1:25 or greater were classified as positive.

*Prediction of the parasite richness depending on the livestock species in the home range*

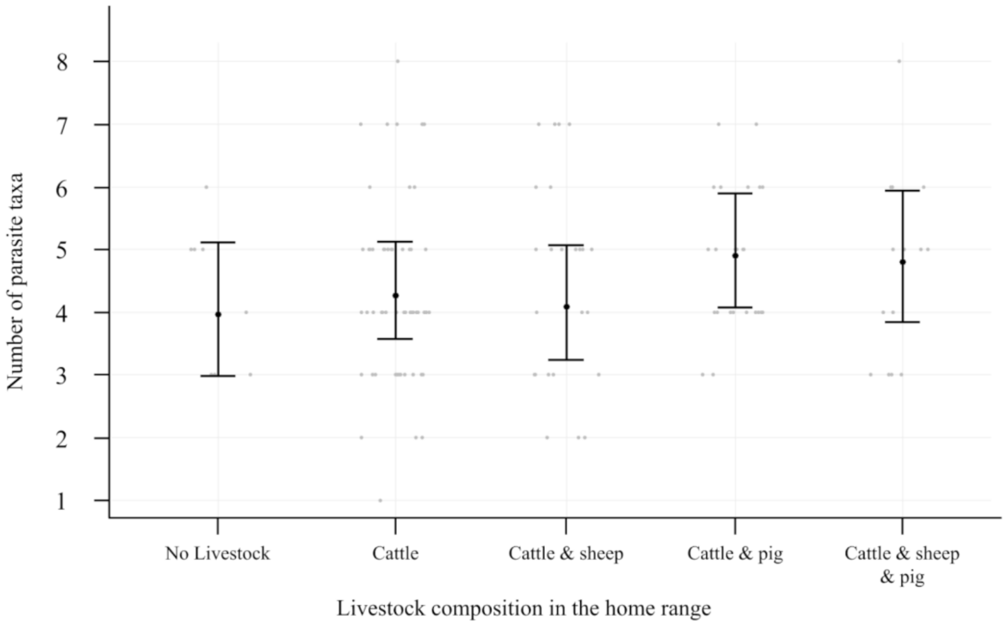


**Figure S1.** Relationship between livestock species present in the home range on the number of parasite taxa predicted by the HMSC model in 130 roe deer.

*Association of host-related variables and parasite mode of transmission*

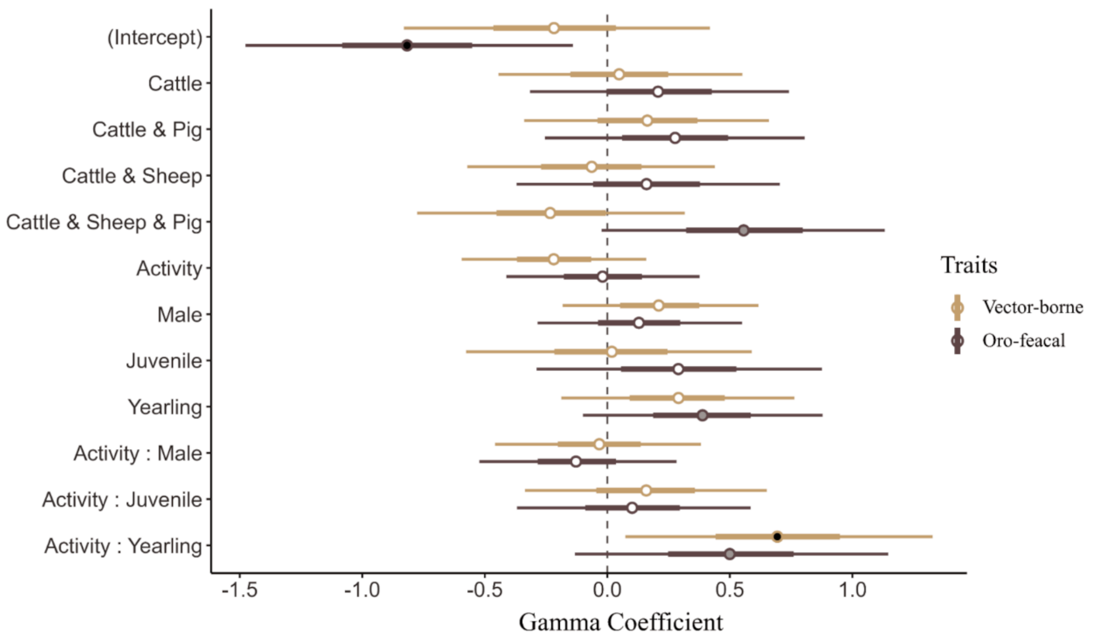


**Figure S2.** Effect of roe deer individual covariates on probability of infection by parasites grouped by transmission mode (faecal-oral or vectorised).
